# CHAS, a deconvolution tool, infers cell type-specific signatures in bulk brain histone acetylation studies of neurological and psychiatric disorders

**DOI:** 10.1101/2021.09.06.459142

**Authors:** Kitty B. Murphy, Yuqian Ye, Maria Tsalenchuk, Alexi Nott, Sarah J. Marzi

## Abstract

Acetylation of histone H3 lysine 27 (H3K27ac) has emerged as an informative disease-associated epigenetic mark. However, cell type-specific contributions to epigenetic dysregulation in disease are unclear as studies have often used bulk brain tissue. Therefore, methods for the deconvolution of bulk H3K27ac profiles are critical. Here we developed the Cell type-specific Histone Acetylation Score (CHAS), a computational tool for inferring cell type-specific signatures in bulk brain H3K27ac profiles. We applied CHAS to > 300 H3K27ac ChIP-seq samples from studies of Alzheimer’s disease, Parkinson’s disease, autism spectrum disorder, schizophrenia, and bipolar disorder in bulk post-mortem brain tissue. In addition to recapitulating known disease-associated shifts in cellular proportions, we identified novel cell type-specific biological insights into brain disorder associated regulatory variation. In most cases, genetic risk and epigenetic dysregulation targeted different cell types, thus suggesting independent mechanisms. For instance, Alzheimer’s disease genetic risk was exclusively enriched within microglia, while epigenetic dysregulation predominantly fell within oligodendrocyte-specific H3K27ac regions. In addition, reanalysis of the original datasets using CHAS enabled identification of biological pathways associated with each neurological and psychiatric disorder at cellular resolution.

## Introduction

H3K27ac is a highly cell type-specific epigenetic modification that marks active enhancers and promoters^1^ and can play a context-dependent, causal role in regulating gene expression ^21^. Brain disorder risk variants predominantly fall into non-coding and regulatory regions^3^, such as those marked by H3K27ac^4^. Given the high cell type specificity and association with transcriptional regulation, integrating genome-wide profiles of H3K27ac from disease-relevant cell types can be useful for functional interpretation of these risk variants. This was demonstrated by efforts mapping regulatory elements to major cell types in the human cortex and investigating neurological and psychiatric disease-risk associations^5^. Additionally, H3K27ac responds to external stimuli^1^, including those associated with disease. Identifying cell type-specific H3K27ac signals in brain disorders can therefore be used to infer dysregulated signalling pathways at cell type resolution.

Studies on post-mortem human brains have identified genome-wide dysregulation of histone acetylation associated with several brain disorders^6–10^. However, interpretation of these studies is limited by the use of bulk tissue, which does not account for the high cellular heterogeneity in the brain. This can lead to biological findings being driven by differences in cellular abundance rather than disease-associated changes, and limits follow-up studies in the appropriate cell types. Thus far, to control for cellular composition, studies have used approaches such as CETS^11^, a tool capable of quantifying neuronal proportions from DNA methylation data, and by measuring the neuronal fraction using flow cytometry. However, these methods require DNA methylation profiles for samples from the same individuals and generally only estimate the proportion of neuronal cell types versus non-neuronal cell types, rather than individual glial cell types.

Although H3K27ac profiling at the purified cell or nuclei population and single-cell level is gaining traction^12,13^, only a few studies have thus far been applied to the human brain^14^, with most only characterising NeuN^+^ and NeuN^-^ samples ^9,10,15^. This highlights an opportunity for the development of cell type deconvolution methods to better interpret bulk brain H3K27ac profiles. Currently, only one approach has been described for the deconvolution of bulk H3K27ac profiles^16^, Here, we present CHAS (Cell type-specific Histone Acetylation Score), a novel computational tool for the cell type deconvolution of bulk brain histone acetylation profiles. To ensure robustness of our model, we implemented two independent algorithms within CHAS and compared their performance across multiple validation approaches and bulk brain histone acetylation studies. The first algorithm, referred to as CHAS, identifies cell type-specific peaks within bulk brain H3K27ac profiles and generates cell type-specific scores which can act as a proxy for cell type proportions. The second algorithm, CHAS-matrix factorisation (MF), is adapted from EPIC, a non-negative matrix factorization model originally designed for the cell type deconvolution of RNA-seq data^17^. We applied both CHAS and CHAS-MF to five brain disorder H3K27ac datasets: Alzheimer’s disease (AD)^7^, Parkinson’s disease (PD)^9^, autism spectrum disorder (ASD)^6^, schizophrenia (SCZ), and bipolar disorder (BPD), to detect shifts in cellular composition and re-investigate differential histone acetylation between cases and controls while controlling for cell type composition. In addition, CHAS allowed identification of cell type-specific genetic risk and epigenetic dysregulation enrichments, which overall highlighted distinct cell types. Thus, CHAS enables deconvolution of H3K27ac in bulk brain tissue, yielding cell type-specific biological insights into brain disease-associated regulatory variation, and is freely available as an R package: https://github.com/Marzi-lab/CHAS.

## Results

### The CHAS model

#### 1. CHAS

Enhancers and H3K27ac domains are known to be highly cell type specific^5,18–20^. CHAS leverages this cell type specificity to annotate peaks identified in bulk brain studies of H3K27ac to their cell type-specific signals in neurons, microglia, oligodendrocytes and astrocytes, as profiled in a previous study^5^. Although we focused on using this reference dataset so that we could perform validation, CHAS could be used with any bulk and reference datasets. CHAS achieves this by overlapping bulk brain H3K27ac peaks with each cell type specific peak set and identifying which of the bulk peaks are specific to a given cell type. For a bulk peak to be defined as cell type-specific two criteria must be met: (i) the bulk peak is annotated only to a single cell type; (ii) the bulk peak overlaps a predefined percentage of that cell type’s peak. This first step in CHAS outputs the bulk peaks annotated to each single cell type, ‘multiple’ cell types (the peak is annotated to more than one cell type), and ‘other’ (the bulk peak is not annotated to any of the reference cell types).

Analysis of bulk tissue can be difficult due to differences in cell type proportion in response to disease, or resulting from discrepancies in brain region sampling. To overcome this, using each set of cell type-specific H3K27ac peaks, CHAS generates cell type-specific scores. By averaging the normalized signal intensity of a sample across all peaks specific to a given cell type, CHAS derives a proxy of the proportion of that cell type in the given bulk sample. For sample y and cell type x, CHAS sums up the peak-normalised counts (s_p,y_; for peak-normalisation see Materials and Methods) across all peaks that are specific to cell type x, divided by the total number of cell type-specific peaks for cell type x. In addition, each score is also normalised by the number of cell type-specific peaks. This normalisation ensures that cell types with a greater number of specific peaks are assigned a higher CHAS score, but do not enable cross-cell type comparisons. As a constraint from peak-normalisation, the maximum signal intensity for any given peak and sample is 1 and any resulting CHAS score will always lie in the interval between 0 and 1.

Of note, given the application of CHAS to deconvolute and control for cellular heterogeneity in histone acetylation studies of brain diseases, we must work under the assumption that disease-related differences in histone acetylation are limited to only a subset of cell type-specific peaks, and that cell-type specific epigenetic variation far outweighs variation associated with disease status^15,21^. We can therefore use cell type-specific chromatin immunoprecipitation sequencing (ChIP-seq) H3K27ac signal intensities to act as a proxy for cell type proportion in bulk tissue data.

Briefly, CHAS requires three inputs:

(i) bulk tissue H3K27ac peaks;
(ii) cell sorted H3K27ac reference peaks;
(iii) counts matrix for the bulk H3K27ac peaks.

CHAS then performs two main analytical tasks (**Fig. 1**):

(i) Identification of cell type-specific peaks in bulk tissue H3K27ac profiles using cell sorted H3K27ac data;
(ii) Generation of cell type-specific scores on the basis of genome-wide average ChIP-seq signal intensities.

**Figure 1.**
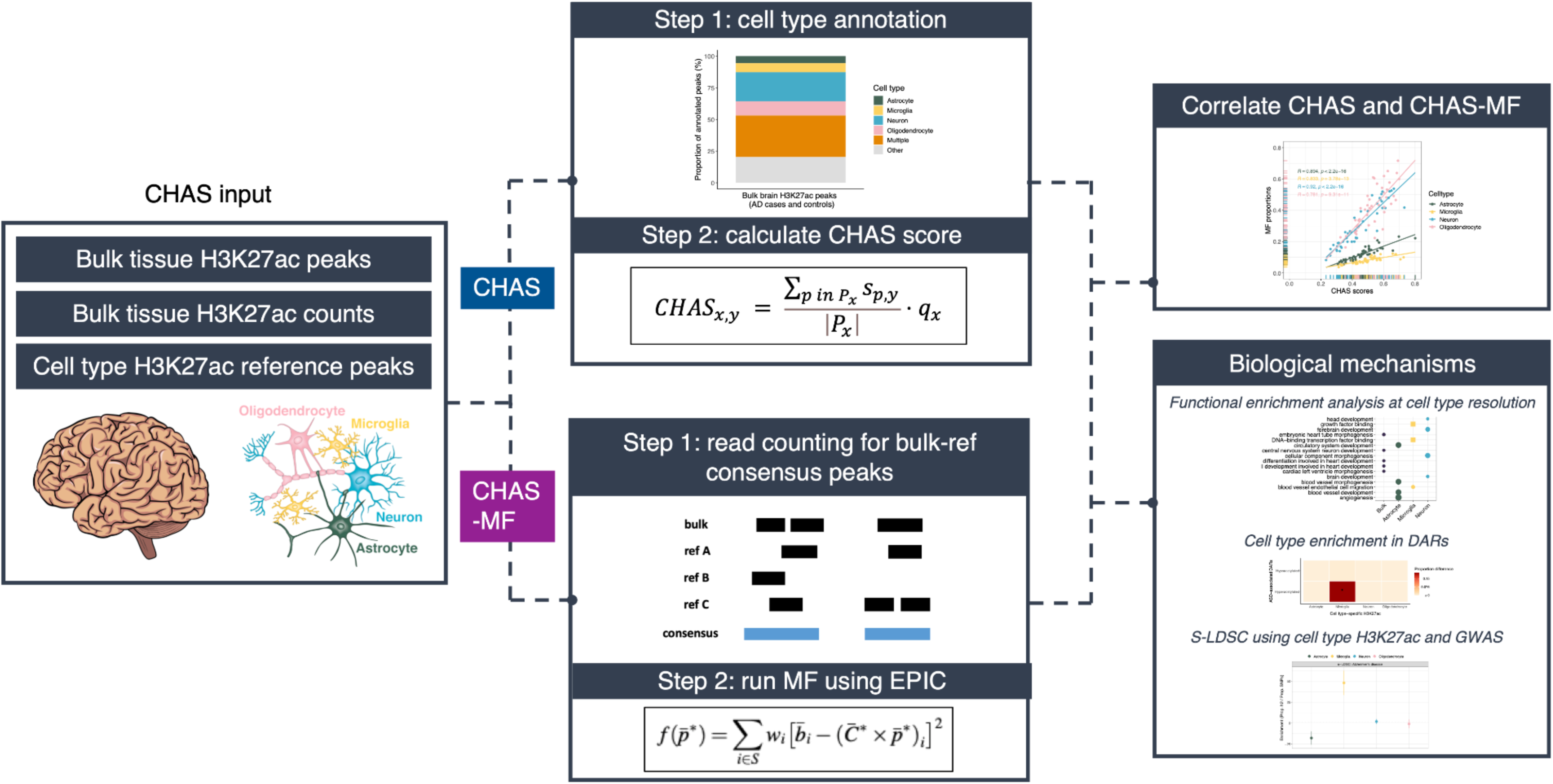
CHAS workflow. CHAS is split into two main computational workflows: CHAS and CHAS-MF. The first step in CHAS is the identification of cell type-specific peaks in bulk tissue H3K27ac profiles, which is achieved using publicly available brain cell sorted H3K27ac data^5^. Using the cell type-specific peaks identified in step 1 and a CPM matrix derived using the bulk tissue H3K27ac data, CHAS generates cell type-specific scores for each sample. Note, for a bulk tissue peak to be defined as cell type-specific, it can only be annotated to a single cell type and it must overlap *n*% of that cell type’s peak(s). The specific required percentage overlap (*n*) can be specified as input to CHAS. The first step in CHAS-MF is merging bulk peaks and reference peaks to create consensus peaks. Each consensus peak is then annotated with corresponding cell types, depending on whether the consensus peak contains cell type-specific peaks. Read counts are then generated for the consensus peaks, either from bam files or existing counts for the bulk and reference samples. Using this count data, CHAS-MF predicts the proportion of each cell type in bulk samples by using EPIC ^22^. The calculation is performed on normalised read counts, controlling for library size and peak length. If multiple reference samples are used for one cell type, the median value will be used as the main input for MF, and the signal variability for each peak is taken into account as weight in MF. The CHAS-derived cell type scores and MF-derived cell type proportions can be correlated, and both can be used as covariates in downstream analyses. *CHAS*𝑥, 𝑦: cell type-specific score for cell type x, sample y; P𝑥: set of cell type-specific peaks for cell type x; S𝑝, 𝑦: standardised peak signal intensity for peak p, sample y. q𝑥: normalisation factor for cell type x. A constraint is applied to S𝑝, 𝑦 whereby for each peak p, the maximum peak signal intensity for any sample equals 1.

The cell type annotation and generation of cell type-specific scores is implemented and freely available in our R package CHAS (https://github.com/Marzi-lab/CHAS).

#### 2. CHAS-MF

We additionally implemented a second algorithm in CHAS for estimating cell type proportions based on non-negative MF. CHAS-MF is based on the EPIC R package^17^, and uses MF to model bulk counts as the sum of cell type-specific counts multiplied by the corresponding cell proportion. CHAS-MF first identifies consensus peaks by merging bulk and reference peaks. Read counts for the consensus peaks are then generated for all the bulk and reference samples using one of two approaches: (i) read counting with bam files if they are available, or (ii) using the bulk and reference counts as a proxy for counts in the consensus peaks.

As EPIC was originally designed for transcriptomic data, we added the three steps to improve its accuracy for deconvoluting bulk brain H3K27ac profiles: (i) bulk and reference H3K27ac counts are normalised based on peak length and library size, (ii) if there are multiple reference samples for a given cell type, the median normalised counts per million (CPM) are used for deconvolution, and (iii) to account for signal variability in each cell type, peaks are assigned weights for matrix factorisation. Higher weights are given to peaks with higher read counts and lower variability. Instead of running MF on all consensus peaks, CHAS-MF selects a set of signature peaks with cell type-specific signals. These are peaks that have high read counts specifically in one cell type with low read counts in all other cell types.

Briefly, CHAS-MF requires three inputs:

(i) bulk tissue H3K27ac peaks;
(ii) cell sorted H3K27ac reference peaks;
(iii) either the bam files for all bulk and reference samples, or the counts matrices for the bulk and reference H3K27ac peaks.

CHAS-MF then performs three main tasks (**Fig. 1**):

(i) Identification of consensus peaks by merging bulk and reference H3K27ac peaks;
(ii) Generation of read counts for consensus peaks in bulk and reference H3K27ac samples;
(iii) Prediction of cell type proportions using MFon the basis of normalised genome-wide ChIP-seq signal intensities.

### Validation of CHAS

To validate the accuracy of CHAS for predicting cell type composition in bulk tissue profiles, we simulated pseudobulk H3K27ac profiles of known cell type composition using the cell sorted H3K27ac data previously described^5^. Each pseudobulk sample was made up of 30 million randomly sampled reads from astrocytes, microglia, neurons, and oligodendrocytes. The cellular composition was based on experimentally quantified proportions of these cell types from the cortex of 17 individuals with AD and 33 age-matched non-AD subjects^23^. We performed peak calling and read count generation for these pseudobulk samples and the resultant peaks and counts were used to run CHAS. For each cell type, we observed a near-perfect correlation between the CHAS score and the true cell type proportion (Pearson’s product moment correlation, R >= 0.99, *p* < 2.2 x 10^-16^ for CHAS and R v>=0.93, *p* < 2.2 x 10^-16^ for CHAS-MF across all cell types, **Fig. 2a,b**). To evaluate the robustness of our approach, this process was repeated with varying read depths (20 and 10 million reads) and sample sizes (25 and 10 samples). We again observed a strong correlation between CHAS derived cell type scores and true cell type proportions (Pearson’s product moment correlation, r >= 0.99) for each analysis; **Supplementary Fig. 1a-d**), highlighting that CHAS is robust to read depth and sample size variation. To further validate our method, we used CHAS and CHAS-MF to deconvolute NeuN^+^ and NeuN^-^ H3K27ac data from two different brain regions of 15 healthy individuals^15^. Across the 15 NeuN^+^ samples from the anterior cingulate cortex (ACC), the mean neuronal proportion when taking into account only cell type-specific peaks was significantly higher than in the 15 NeuN^-^ samples from the same individuals (CHAS, Welch’s t-test, mean difference = 0.86, p = 4.35 x 10^-14^ ; CHAS-MF, paired t-test, mean difference = 0.85, *p* < 2.2 x 10^-16^). The same was observed for the 14 NeuN^+^ and 14 NeuN^-^ samples from the dorsolateral prefrontal cortex (PFC) (CHAS, paired t-test, mean difference = 0.89, p < 2.2^-16^; CHAS-MF, paired t-test, mean difference = 0.90, *p* < 2.2 x 10^-16^).

**Figure 2.**
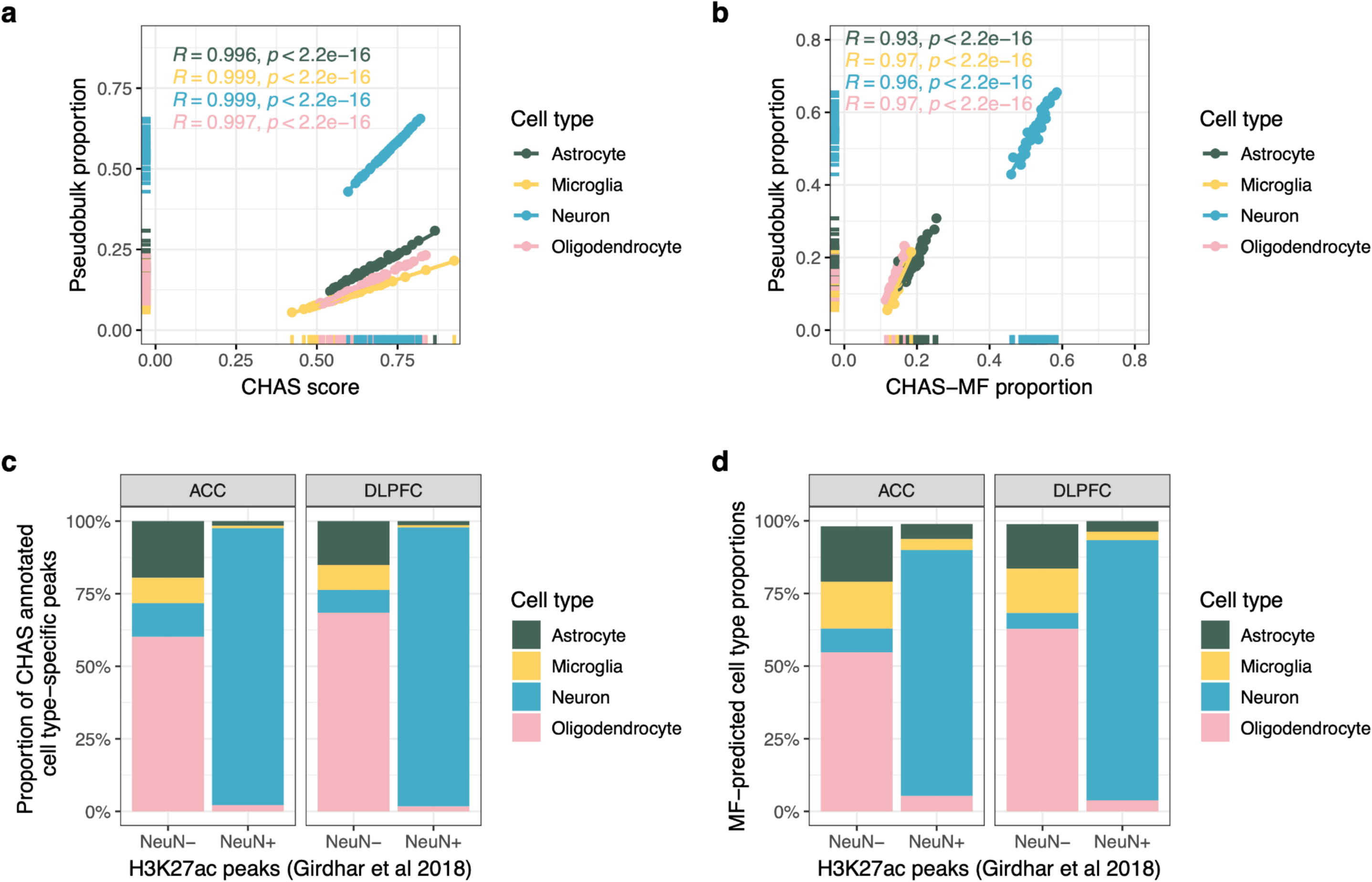
CHAS cell type scores and proportions correlate with true cell type proportions. **a** Scatterplot ofthe CHAS-derived cell type score against the true proportion of the cell type in the pseudobulk sample. **b** Scatterplot of the CHAS-MF proportions against the true proportion of the cell type in the pseudobulk sample. **c** Proportion of CHAS-annotated cell type-specific peaks within NeuN^-^ and NeuN^+^ H3K27ac profiles. **d** CHAS-MF derived cell type proportions within NeuN^-^ and NeuN^+^ H3K27ac profiles.

### Deconvolution of bulk brain H3K27ac in Alzheimer’s disease highlights oligodendrocyte-specific epigenetic dysregulation

AD is a complex, highly heritable neurodegenerative disorder. Genetic variants contributing to AD susceptibility predominantly fall into non-coding and regulatory regions^3^, such as those marked by H3K27ac ^4,5^. Aberrations in H3K27ac associated with AD have been reported in the human brain^7,8^, however, the contribution of individual cell types in epigenetic dysregulation associated with the disease is only beginning to be dissected ^14^ To address this, we used CHAS to deconvolute H3K27ac profiles from the entorhinal cortex of 24 AD cases and 23 controls^7^. Briefly, Marzi et al. (2018) found widespread dysregulation of H3K27ac associated with AD, with differentially acetylated regions (DARs) identified in the vicinity of known early-onset AD risk genes, as well as genomic regions containing variants associated with late-onset AD.

In this study, we re-analysed the original data using CHAS. Out of 183,353 peaks, 80% (n = 146,144) were annotatable to one or more cell types, with 47% (n = 85,824) specific to a single cell type (**Fig. 3a**). Using CHAS-MF, we then estimated cell types proportions in each bulk sample, and found that the four cell types constituted on average 92% of the samples (**Fig. 3b**). In the original study, neuronal proportion (NeuN⁺ fraction) estimates for samples from the same individuals had been derived based on matched bulk brain DNA methylation data using CETS^7,11^, a tool for estimating neuronal proportion. CHAS-derived neuronal scores and CETS-derived neuronal proportion estimates correlated across the 47 samples (Spearman’s rank correlation coefficient, r = 0.409, *p* = 0.00429, **Supplementary Fig. 1e**).

**Figure 3:**
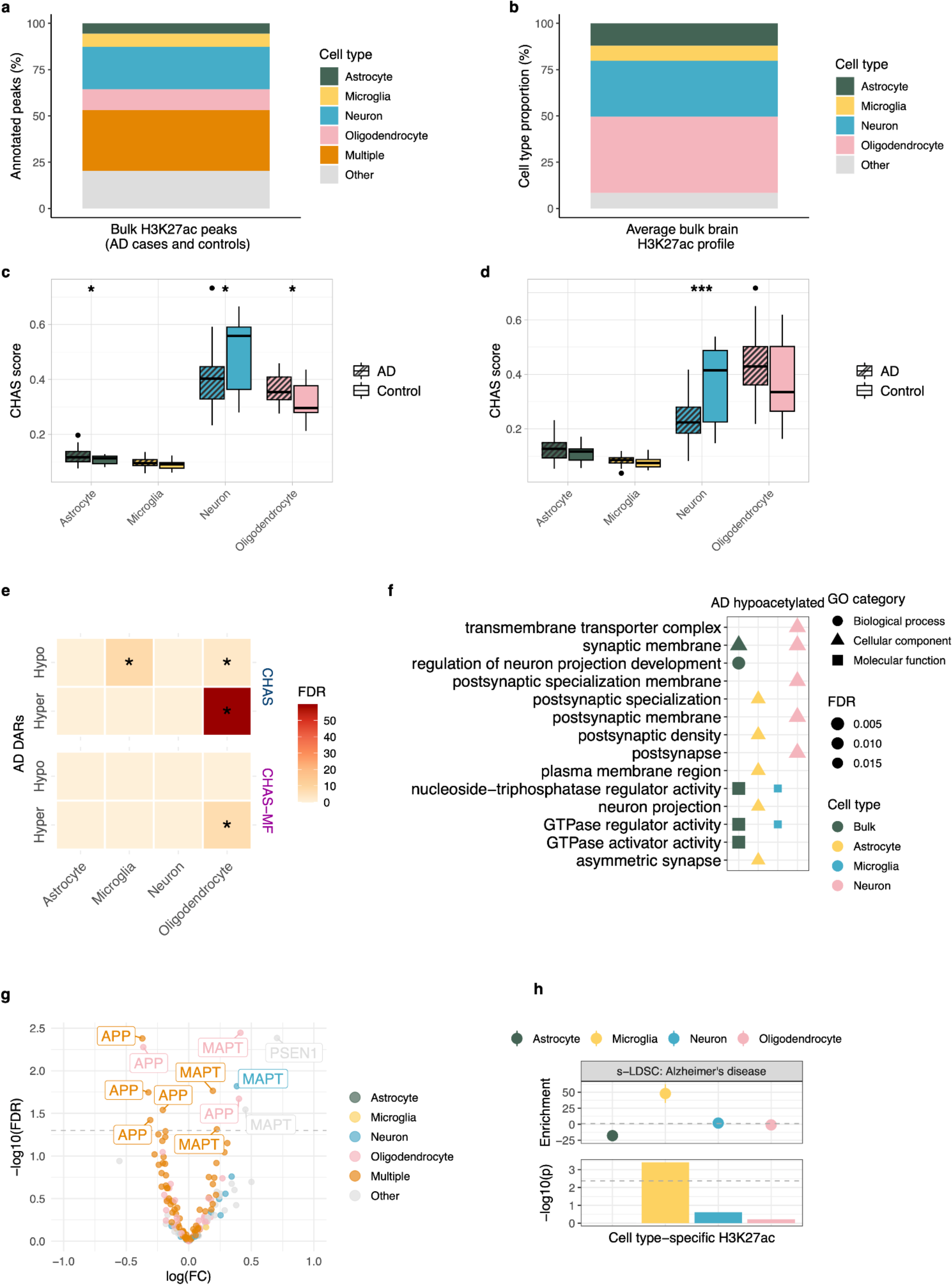
Cell type deconvolution of H3K27ac profiles from the AD brain highlights a role for oligodendrocytes and microglia. **a** Stacked barplot of proportion of bulk brain H3K27ac peaks (AD cases and controls) annotated to astrocytes, microglia, neurons, oligodendrocytes, multiple (at least two cell types), or other (none of the reference cell type peaks) using CHAS. **b** Stacked barplot of average cell type proportions across all bulk brain H3K27ac profiles from AD cases and controls, estimated using CHAS-MF. **c** Boxplot comparing the CHAS cell type score means between AD cases and controls. The neuronal score was significantly lower in AD cases when compared to controls, whereas the astrocyte and oligodendrocyte score was higher. P values were calculated using Welch’s two-sample t-test. **d** Boxplot comparing the MF cell type proportions between AD cases and controls. The neuronal proportion was significantly lower in AD cases when compared to controls. P values were calculated using Welch’s two-sample t-test. **e** Heatmap of cell type enrichments amongst AD-associated DARs identified when controlling for CHAS scores and CHAS-MF proportions, calculated using the hypergeometric test. Hypergeometric p values were corrected using FDR. **f** Functional enrichment analysis using bulk brain and cell type-specific AD-associated hyperacetylated regions controlling for CHAS scores. Shown are the top 10 enriched pathways for each cell type. P values were corrected using FDR. **g** Volcano plot of all H3K27ac peaks annotated to the AD genes APP, PSEN1, PSEN2, and MAPT. Labelled peaks are those that were differentially acetylated between AD cases and controls. **i** Results from s-LDSC using an AD GWAS^35^ and CHAS-annotated cell type-specific H3K27ac peaks from the brains of individuals with AD, and controls. In the top panel, the y-axis is the enrichment value output by LDSC. In the bottom panel, the y-axis represents the coefficient *p* value transformed from the coefficient z-score output by LDSC. The grey dashed line at -log10(P) = 2.4 is the cutoff for Bonferroni significance. AD SNP heritability was found to be exclusively enriched in microglia-specific H3K27ac peaks.

We next used CHAS to evaluate shifts in cellular composition in the bulk brain data, testing whether these replicate known disease-associated changes. To this end, we compared CHAS-derived scores and MF-derived proportions between AD cases and controls for each cell type. As expected, given that AD is primarily associated with neuronal loss, we observed a lower neuronal score in AD brains compared to controls (Welch’s t-test, two-sided, difference in mean score = 0.08, *p* = 0.028; **Fig. 3c**). The same was observed when comparing the MF-derived neuronal proportion in AD cases vs controls (Welch’s t-test, two-sided, difference in mean proportion = 0.13, *p* = 0.001; **Fig. 3d**). We also report a higher oligodendrocyte score (Welch’s t-test, two-sided, difference in mean score = 0.07, *p* = 0.013; **Fig. 3c**) but not proportion in AD cases (Welch’s t-test, two-sided, difference in mean proportion = 0.06, *p* = 0.15; **Fig. 3d**). Neither microglia score nor proportion were significantly different between AD cases and controls (Welch’s t-test, two-sided, difference in mean score = 0.05, *p* = 0.072; difference in mean proportion = 0.008, *p* = 0.21; **Fig. 3c-d**).

We then used the cell type scores and proportions to re-investigate differential histone acetylation in AD at cell type resolution. Employing the quasi-likelihood *F* test in edgeR^24^ we first quantified differential acetylation between AD cases and controls, while controlling for cell type scores (CHAS in neurons, astrocytes, microglia and oligodendrocytes) and age at death. 5,763 peaks were characterised by AD-associated hyperacetylation and 5,904 were characterised by AD-associated hypoacetylation (false discovery rate (FDR) < 0.05) (**Supplementary Table 1**). To evaluate the likelihood of false-positive associations, we repeated the differential histone acetylation analysis using permuted AD case and control labels. Across 100 permuted datasets there was never more than one significant peak at FDR < 0.05, thus making it unlikely that the results of our differential acetylation analysis based on the true AD case and control labels were detected due to chance. Differential acetylation analysis was repeated using MF-derived proportions for the four cell types and age at death as covariates. In contrast to controlling for CHAS scores, this analysis revealed a significantly lower number of AD-associated DARs to the original study^7^, with 324 hyper- and 792 hypo-acetylated peaks (**Supplementary Table 1**). Notably, AD-associated hyperacetylated regions from both analyses (controlling for CHAS scores versus proportions) were significantly enriched for oligodendrocytes when compared to regions that were not differentially acetylated (hypergeometric test, *p* = 4.33 x 10^-57^ when controlling for scores, *p* = 3.25 x 10^-8^ when controlling for proportions; **Fig. 3e**). A similar enrichment of oligodendrocyte- and microglia-specific peaks was observed for AD-associated hypoacetylated regions when controlling for cell type scores (hypergeometric test *p* = 8.87 x 10^-3^ for oligodendrocytes; *p* = 2.48 x 10^-5^ for microglia; **Fig. 3e**). These results are consistent with a previous study in which oligodendrocytes were reported to show the most widespread acetylation differences in the AD brain^25^. In addition, the top AD-associated hyperacetylated peak when controlling for CHAS scores was specific to oligodendrocytes and located in the vicinity of *MVB12B*, a gene implicated in vesicular trafficking (**Supplementary Table 1**). This peak was also differentially acetylated when controlling for cell type proportions **(Supplementary Table 1)**. *MVB12B* has previously been identified as an AD risk gene^26^ and forms part of an oligodendrocyte-enriched gene network in the AD brain^27^. One of the top-ranked AD-hypoacetylated peaks when controlling for CHAS scores was annotated to multiple cell types, and located near *POC1B*, an AD risk gene that was reported to form part of the same core oligodendrocyte gene network as *MVB12B*^27^. This peak was also differentially acetylated when controlling for cell type proportions **(Supplementary Table 2)**. To provide a broader overview of the concordance between the two analyses, we compared the logFC values for acetylation and found a strong correlation (**Supplementary Fig. 2a**).

Using clusterProfiler^28^, we were able to match the functional categories associated with AD differentially acetylated bulk peaks to their cell types (**Fig. 3f, Supplementary Fig. 3**). Additionally, we were able to identify cell type-specific dysregulated pathways that were not seen in functional enrichment analyses based on bulk peaks. Neuron-specific hyperacetylated peaks were enriched for synaptic functions, especially in the postsynaptic membrane (**Fig. 3f**). This suggests an active adaptation of synaptic density and functions in the AD brain, which has been reported by several studies^29–31^. On the other hand, microglia-specific hypoacetylated regions were enriched for GTPase activity, which is important for homeostatic functions in microglia that regulate neuronal physiology^32,33^ (**Fig. 3g**). As reported previously^7^, differential H3K27ac was observed in regulatory regions annotated to genes *MAPT*, *APP*, *PSEN1,* and *PSEN2*, which are known to be associated with early onset AD or directly involved in AD-neuropathology (**Supplementary Table 1**). To link CHAS-annotated cell type specific acetylation with genetic risk for AD, we performed partitioned heritability analysis^34^ and quantified enrichment of AD GWAS risk variants within cell type-specific H3K27ac regions annotated in the bulk data set. As reported previously, significant enrichment of AD risk loci was found within microglia-specific H3K27ac regions but not in the other cell types^5^ (**Fig. 3h**).

### Cortical H3K27ac patterns in the Parkinson’s disease brain are associated with oligodendrocytes

The role of H3K27ac at the cellular level in PD remains largely unexplored. Although cell type vulnerability in PD is commonly attributed to dopaminergic neurons, genetic risk has been variably associated with cholinergic and enteric neurons, as well as oligodendrocytes^36^. Furthermore, cortical epigenetic studies have thus far failed to yield significant cell type enrichments. However, as our deconvolution of epigenetic signatures of AD highlights, genetic risk and epigenetic dysregulation are not necessarily in agreement in relation to cell types. Therefore, we applied CHAS to a bulk brain H3K27ac study in PD cases and controls. Toker et al. (2022) observed genome-wide dysregulation of histone acetylation in PFC of individuals with PD from two independent cohorts: the ParkWest (PW) study cohort^37^ and the Netherlands Brain Bank (NBB) cohort (https://www.brainbank.nl/). The authors reported that PD-associated hyperacetylated regions were annotated to genes implicated in PD pathology, and also describe decoupling between promoter H3K27ac and gene expression in the PD brain^9^. To account for cellular heterogeneity they integrated H3K27ac regions differing between NeuN⁺ and NeuN⁻ cell types with brain cell type-specific marker genes, and employed a principal component analysis approach to serve as a proxy for cell type composition across their samples^9^. Using this approach, they found no significant differences in cell type proportion between PD cases and controls.

Using the PW cohort (13 PD cases and 10 controls) peaks and read counts generated by Toker et al. (2022)^9^, we filtered out peaks with low read counts and peaks annotated to non-canonical chromosomes before running CHAS. Out of 132,340 peaks, 74% were annotatable to at least one cell type in the PW cohort (**Fig. 4a**). Using CHAS-MF, we estimated the cell type proportion in each bulk sample, and found that the four cell types constituted on average 96% of the samples (**Fig. 4b**). Cell proportions estimated by CHAS-MF from PD and control H3K27ac samples correlated with CHAS scores in all four cell types (**Supplementary Fig 3b**). In line with the original study, we found no significant difference in cell type scores or proportions between PD cases and controls (**Fig. 4c-d**). While this indicates that bulk PFC tissue may be less prone to confounding disease-associated shifts in cellular proportions in PD, it simultaneously does not represent the primarily disease-affected brain region. Differential histone acetylation analysis controlling for sex, age, and CHAS scores identified four hyperacetylated and three hypoacetylated peaks, of which four were oligodendrocyte-specific and one microglia-specific (**Fig. 4e**). Repeating the analysis using MF proportions instead of CHAS scores revealed 43 peaks characterised by hyperacetylation and 21 peaks characterised by hypoacetylation, and similarly, a large proportion of these peaks were annotated to oligodendrocytes (**Fig. 4e; Supplementary Table 2**). Common genes were found to be in the vicinity of DARs while controlling for CHAS and CHAS-MF respectively, (**Supplementary Table 2**), and logFC values of acetylation between the two analyses were strongly correlated (**Supplementary Fig. 2b**). Due to the low number of DARs, we chose not to perform cell type- or functional enrichment analysis. We also performed partitioned heritability analysis to quantify enrichment of PD GWAS risk variants within cell type-specific H3K27ac regions, but did not observe significant enrichment of PD risk in any of the cell type specific H3K27ac peak sets (**Fig. 4f**).

**Figure 4:**
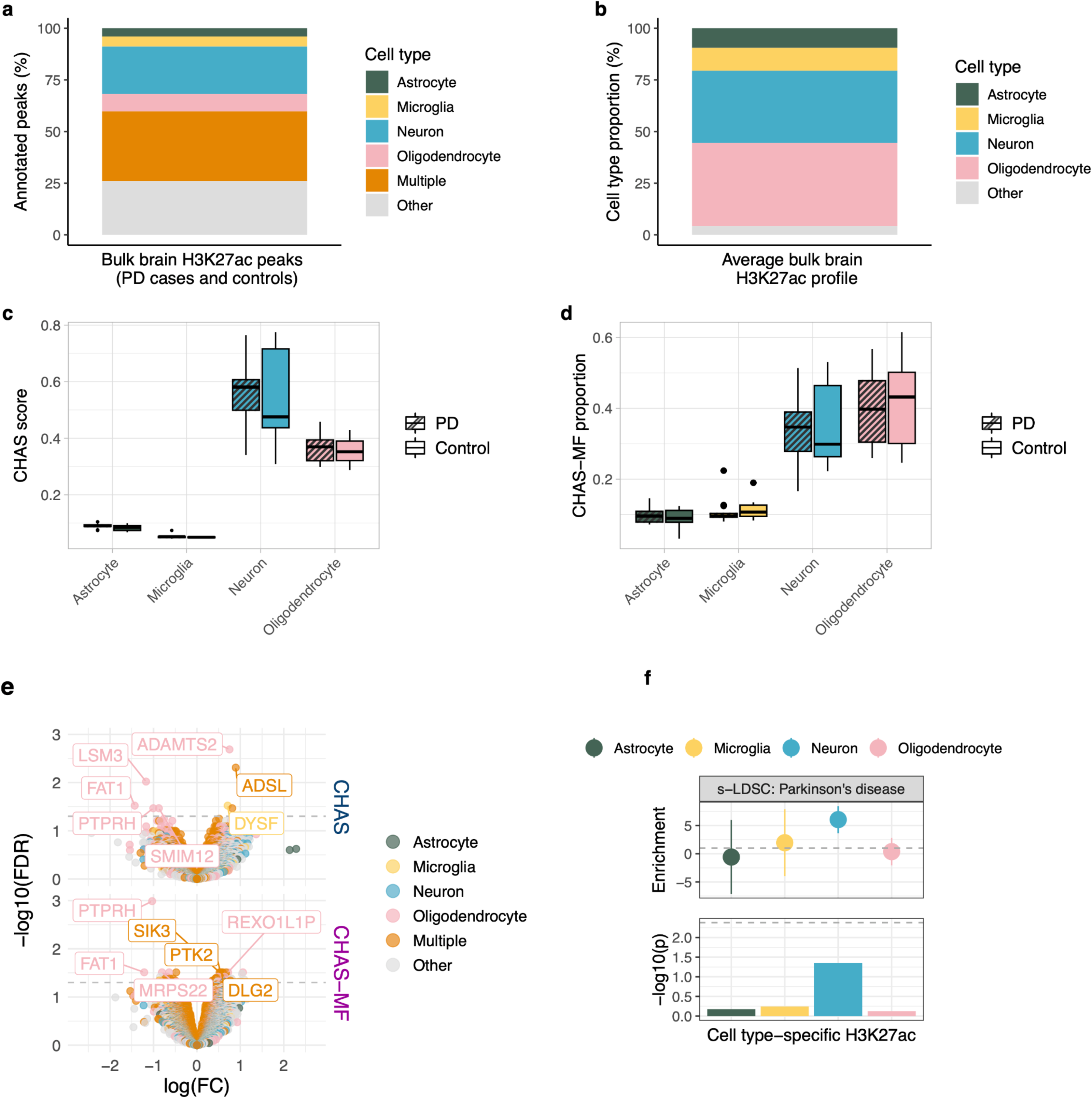
Cortical H3K27ac patterns in the PD brain are associated with oligodendrocytes. **a** Stacked barplot of proportion of bulk brain H3K27ac peaks (PD cases and controls) annotated to astrocytes, microglia, neurons, oligodendrocytes, multiple (at least two cell types), or other (none of the reference cell type peaks) using CHAS. **b** Stacked barplot of average cell type proportions across all bulk brain H3K27ac profiles from PD cases and controls, estimated using CHAS-MF. **c** Boxplot comparing the CHAS cell type score means between PD cases and controls. No significant differences were observed. P values were calculated using Welch’s two-sample t-test. **d** Boxplot comparing the MF cell type proportions between PD cases and controls, no significant differences were observed. P values were calculated using Welch’s two-sample t-test. **e** Volcano plot of differentially acetylated regions between PD cases and controls when controlling for age, sex, and either CHAS scores or CHAS-MF proportions. Peaks annotated to genes are those with FDR < 0.05. **f** Results from s-LDSC using a PD GWAS ^38^ and CHAS-annotated cell type-specific H3K27ac peaks from the brains of individuals with PD, and controls. In the top panel, the y-axis is the enrichment value output by LDSC. In the bottom panel, the y-axis represents the coefficient p value transformed from the coefficient z-score output by LDSC. The grey dashed line at -log10(P) = 2.4 is the cutoff for Bonferroni significance.

### Cell type deconvolution of autism-associated H3K27ac highlights an epigenetic role for microglia

ASD encompasses a group of genetically complex and heterogeneous neurodevelopmental disorders. At the bulk tissue level, dysregulation of H3K27ac in ASD brains is associated with genes involved in synaptic transmission and immunity, as well as genes which harbour rare ASD mutations^6^. Taking into consideration the high cell type-specificity of H3K27ac, CHAS provides an opportunity to re-investigate these epigenomic perturbations at cell type resolution. Previously, Sun et al. (2016)^6^ performed a histone acetylome-wide association study across three different brain regions from ASD cases and age-matched controls. They reported widespread dysregulation of H3K27ac in PFC and temporal cortex of ASD cases, with similar changes observed in both brain regions. In contrast, only a small proportion of peaks were differentially acetylated in cerebellum^6^.

Using 80 ChIP-seq samples from PFC (40 cases, 40 controls), and 62 samples from cerebellum (31 cases, 31 controls), we called peaks in each brain region using MACS2^39^. After filtering out peaks with low read counts, we defined an optimal peak set for each brain region: 246,300 peaks in PFC, and 236,358 peaks in cerebellum. We then used these optimal peak sets to run CHAS to evaluate cell type proportions in each brain region, and to generate cell type-specific scores and MF-derived proportions for each sample in each brain region. In the PFC, 73% of bulk peaks were annotatable to at least one cell type, whereas in the cerebellum 49% of peaks could be annotated to a cell type (**Fig. 5a**). This is most likely explained by epigenetic differences across brain regions in the reference and test datasets^40,41^: brain region specific differences could exist in the epigenetic state of the same cell type, for instance microglia across multiple brain regions. Similarly, there can be differences in the actual cell types located in different brain regions. For example, the cortex contains highly specialised pyramidal neurons, while purkinje neurons are specific to the cerebellum. Using CHAS-MF, we estimated the cell type proportion in each bulk sample, and found that the four cell types constituted on average 95% of the PFC samples and 92% of the cerebellum samples (**Fig. 5c**).

**Figure 5:**
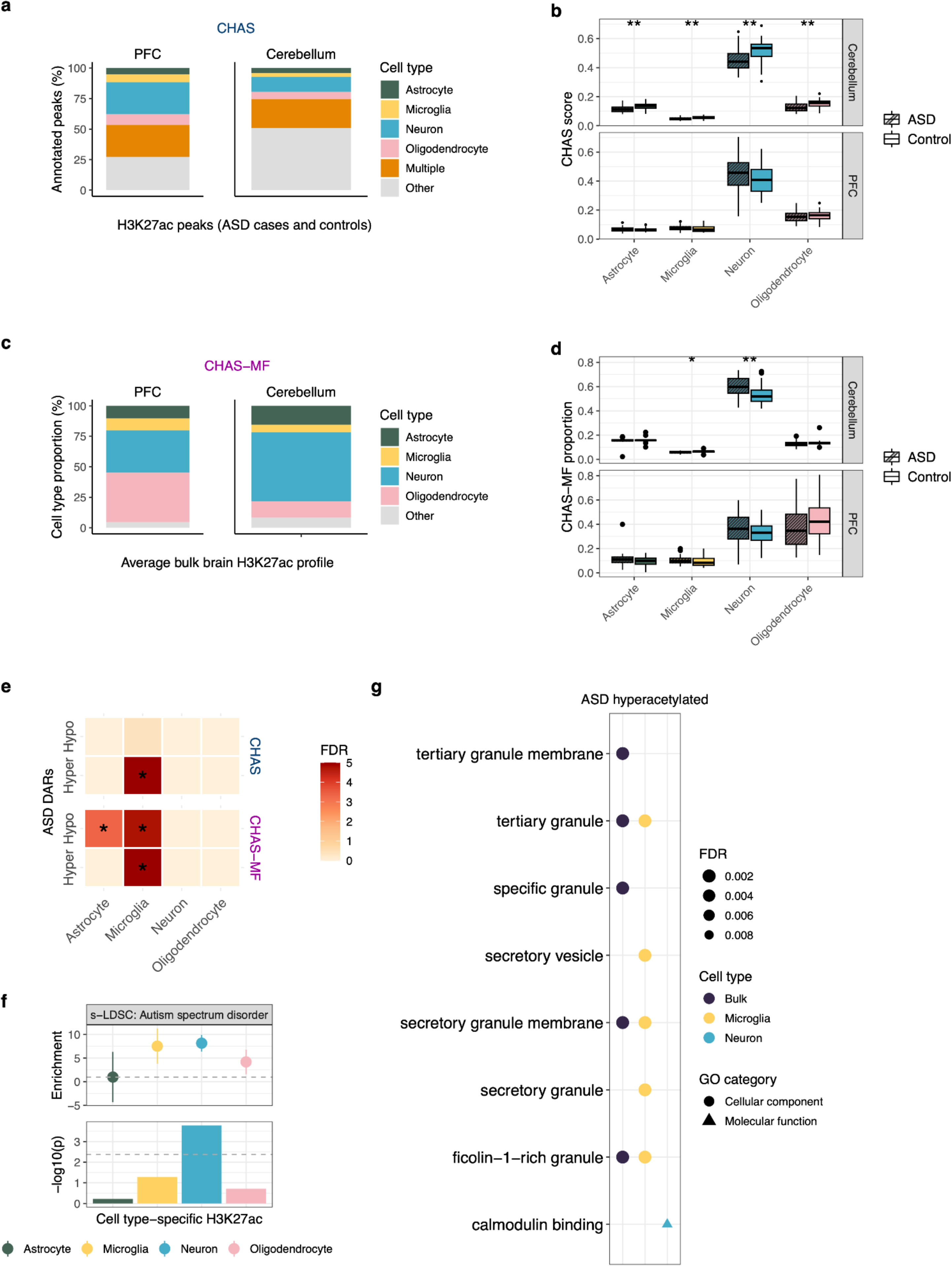
Cell type deconvolution of autism-associated H3K27ac highlights a role for microglia. **a** Stacked barplot of proportion of bulk PFC and cerebellar H3K27ac peaks (ASD cases and controls) annotated to astrocytes, microglia, neurons, oligodendrocytes, multiple (at least two cell types), or other (none of the reference cell type peaks) using CHAS. **b** Stacked barplot of average cell type proportions across all bulk PFC and cerebellar H3K27ac profiles from ASD cases and controls, estimated using CHAS-MF. **c** Boxplot comparing the CHAS cell type score means between ASD cases and controls. No significant differences were observed for the PFC, whereas all cell type scores were significantly lower in the cerebellum of ASD cases when compared to controls. P values were calculated using Welch’s two-sample t-test. **d** Boxplot comparing the MF cell type proportions between ASD cases and controls. P values were calculated using Welch’s two-sample t-test. **e** Heatmap of cell type enrichments amongst ASD-associated DARs identified when controlling for CHAS scores and CHAS-MF proportions,, calculated using the hypergeometric test. Hypergeometric p values were corrected using FDR. **f** Results from s-LDSC using an ASD GWAS and CHAS-annotated cell type-specific H3K27ac peaks from the brains of individuals with ASD and controls. In the top panel, the y-axis is the enrichment value output by LDSC. In the bottom panel, the y-axis represents the coefficient p value transformed from the coefficient z-score output by LDSC. The grey dashed line at -log10(P) = 2.4 is the cutoff for Bonferroni significance. ASD SNP heritability was found to be exclusively enriched in neuron-specific H3K27ac peaks. **g** Functional enrichment analysis using bulk brain and cell type-specific ASD-associated hyperacetylated regions. Shown are the top eight enriched pathways for each cell type. *P* values were corrected using the FDR.

Next, we compared the cell type scores and proportions in ASD cases with controls, and found no significant difference in neither the cell type scores nor the estimated proportions (**Fig. 5b,d**). In cerebellum, we obtained significantly lower cell type scores for all four cell types in ASD cases when compared to controls (**Fig. 5b**). This likely relates to cellular composition differences between cortex and cerebellum, highlighting the need for a brain region appropriate reference. In contrast, deconvolution using CHAS-MF revealed a significant difference exclusively in the neuronal proportion, which might be partly due to the low abundance and high variability of the glial populations (**Fig 5d**). Taken together, the observed ASD-associated differences in cell type scores and proportions in the cerebellum should be interpreted with caution. Based on the cell type annotation of cerebellar peaks, we decided not to perform downstream analyses on this dataset.

Differential histone acetylation analysis controlling for sex, age at death and cell type score or cell type proportion revealed ASD-associated DARs in PFC (7,652 DARs when controlling for scores, 5,320 DARs when controlling for proportions; **Supplementary Table 3**). ASD-associated hyperacetylated regions in PFC, when controlling for cell type scores and proportions, were significantly enriched for microglia when compared to the background peak set (hypergeometric test, *p* < 2.2^-16^; **Fig. 5e**). In addition, cell type enrichment while controlling for cell type proportions revealed astrocyte- and microglia-specific enrichment in ASD-associated hypoacetylated regions (hypergeometric test, *p* = 6.0 x 10^-4^ for astrocyte and *p* = 1.8 x 10^-4^ for microglia; **Fig. 5e**). This is consistent with existing evidence that ASD patients have altered microglial states^42,43^. In addition, the top ranking microglia-specific ASD-associated hyperacetylated peak was located ∼46kb upstream of *EMSY* (**Supplementary Table 3)**. Transcriptome analysis of cortical samples from ASD cases and controls identified *EMSY* as one of the top differentially expressed genes and suggested a general role for dysregulated microglial genes^44^. Moreover, whole exome sequencing in individuals with ASD revealed a de novo loss-of-function mutation in this gene^45^. The top ranking neuron-specific ASD-associated hypoacetylated peak was located 330bp upstream of *COBLL1*, and was identified among the top five differentially acetylated regions while controlling for both CHAS scores and MF proportions. *COBLL1* was recently identified in a quantitative genome-wide association study using MAGMA to be associated with joint attention and nonverbal communication in ASD patients^46^. As observed for the previous datasets, comparison of the logFC values of acetylation between the two analyses revealed a strong correlation (**Supplementary Fig. 2c**). Furthermore, we quantified enrichment of ASD GWAS risk variants within cell type-specific H3K27ac. As reported previously^5^, significant enrichment of ASD risk loci was found within neuron-specific H3K27ac regions (**Fig. 5f**).

Performing functional enrichment analysis at cell type resolution allowed us to overlap the categories associated with the differentially acetylated bulk peaks, with the cell types they were specific to, as well as identify distinct functional enrichments across the four cell types that were not identified in the bulk analysis. For example, as reported in the original study^6^, hyperacetylated peaks in the PFC while controlling for CHAS scores were associated with calmodulin binding (**Fig. 5g; Supplementary Table 3**). Using CHAS, we additionally report that this enrichment is driven by hyperacetylated peaks that were annotated to neurons (**Fig. 5g; Supplementary Table 3**). In addition, ASD hyperacetylated bulk peaks revealed enrichment for immune related processes, which were predominantly specific to microglia (**Fig. 5g; Supplementary Table 3**). This is in line with the growing body of evidence highlighting the role of glial cells and neuroimmune alterations in ASD^42,44,47–49^. Additional bulk and cell type-specific functional enrichments can be seen in **Supplementary Fig. 5**.

### Deconvolution of bulk PFC H3K27ac in schizophrenia and bipolar disorder reveals neuron- and oligodendrocyte-specific epigenetic dysregulation

Until recently, the epigenomic landscapes of schizophrenia and bipolar disorder remained largely unknown. A landmark study in which 249 bulk brain (PFC) samples (133 controls, 68 schizophrenia cases, and 48 bipolar disorder cases) were profiled for H3K27ac reported genome-wide alterations of this epigenetic mark across the disorders, both at the tissue and neuronal level^10^. Although the study accounted for cell type heterogeneity in the bulk brain samples by controlling for the estimated proportions of oligodendrocytes, glutamatergic neurons, and GABAergic neurons, there is added value in re-investigating this dataset additionally using cell type-specific signals in astrocytes and microglia.

Publicly available FASTQ files for all 249 PFC samples were downloaded and pre-processed, before mapping to GRCh38 using Bowtie2^50^. Peak calling was performed using MACS2^39^ on a merged file of all samples. The bulk PFC H3K27ac peaks were annotated to their cell type-specific signals, and cell type scores and proportions were estimated using CHAS and CHAS-MF, respectively. Despite both the bulk and the cell type reference H3K27ac datasets being from the cortex, only 55% of bulk peaks were annotatable to at least one cell type (**Fig 6a**). This could be explained by technical issues, for instance, the cell type reference samples were from fresh resected tissue whereas the bulk profiles were from the post-mortem brain where sample quality can be further affected by post-mortem interval. The smaller than expected overlap could also be explained by biological differences; the reference data are from young brains with epilepsy whereas the bulk data brains are from a whole spectrum of ages and include both healthy and diseased brains. Using CHAS-MF, we estimated the cell type proportion in each bulk sample, and found that the four cell types constituted on average 92% of the samples (**Fig. 6b**).

**Figure 6:**
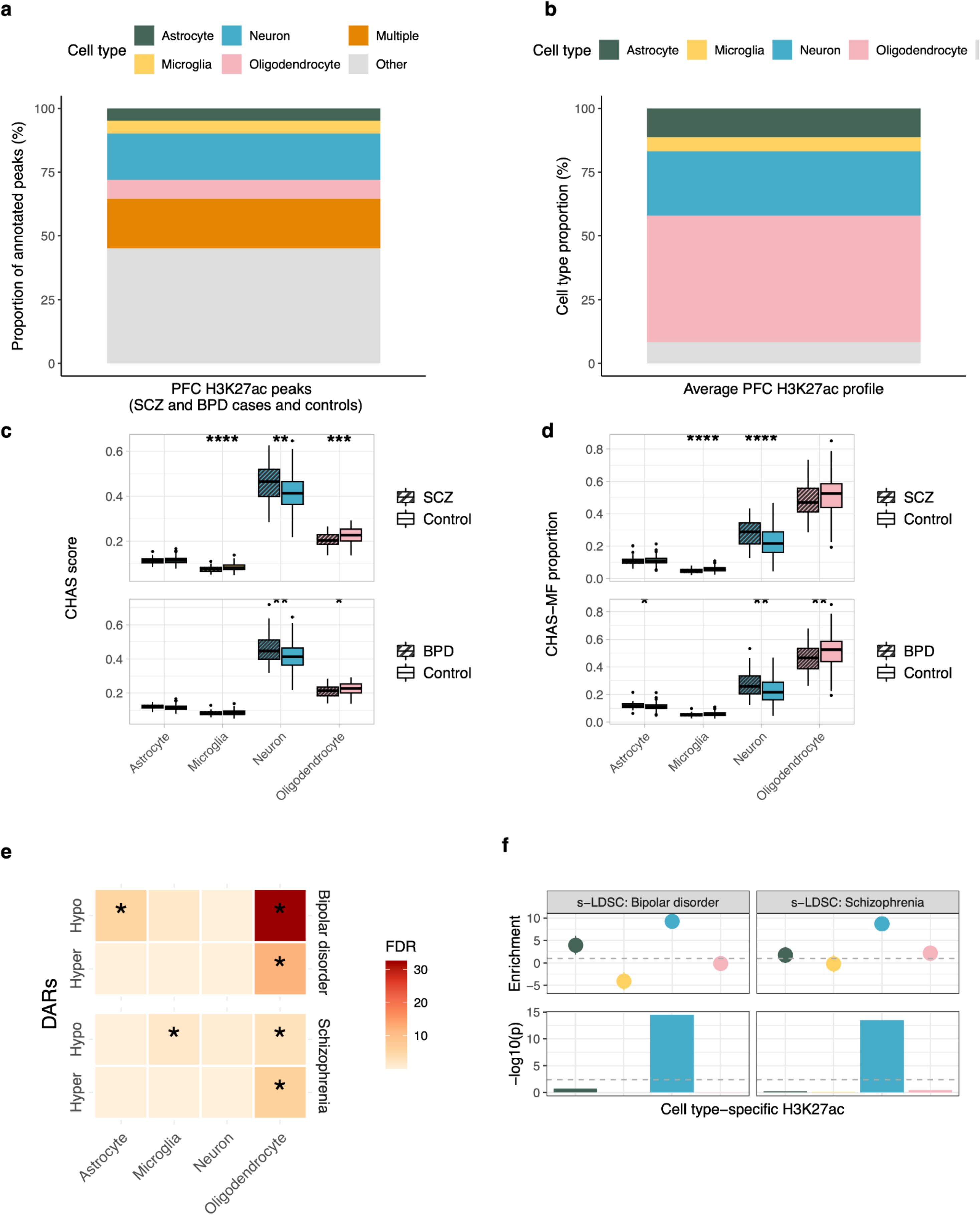
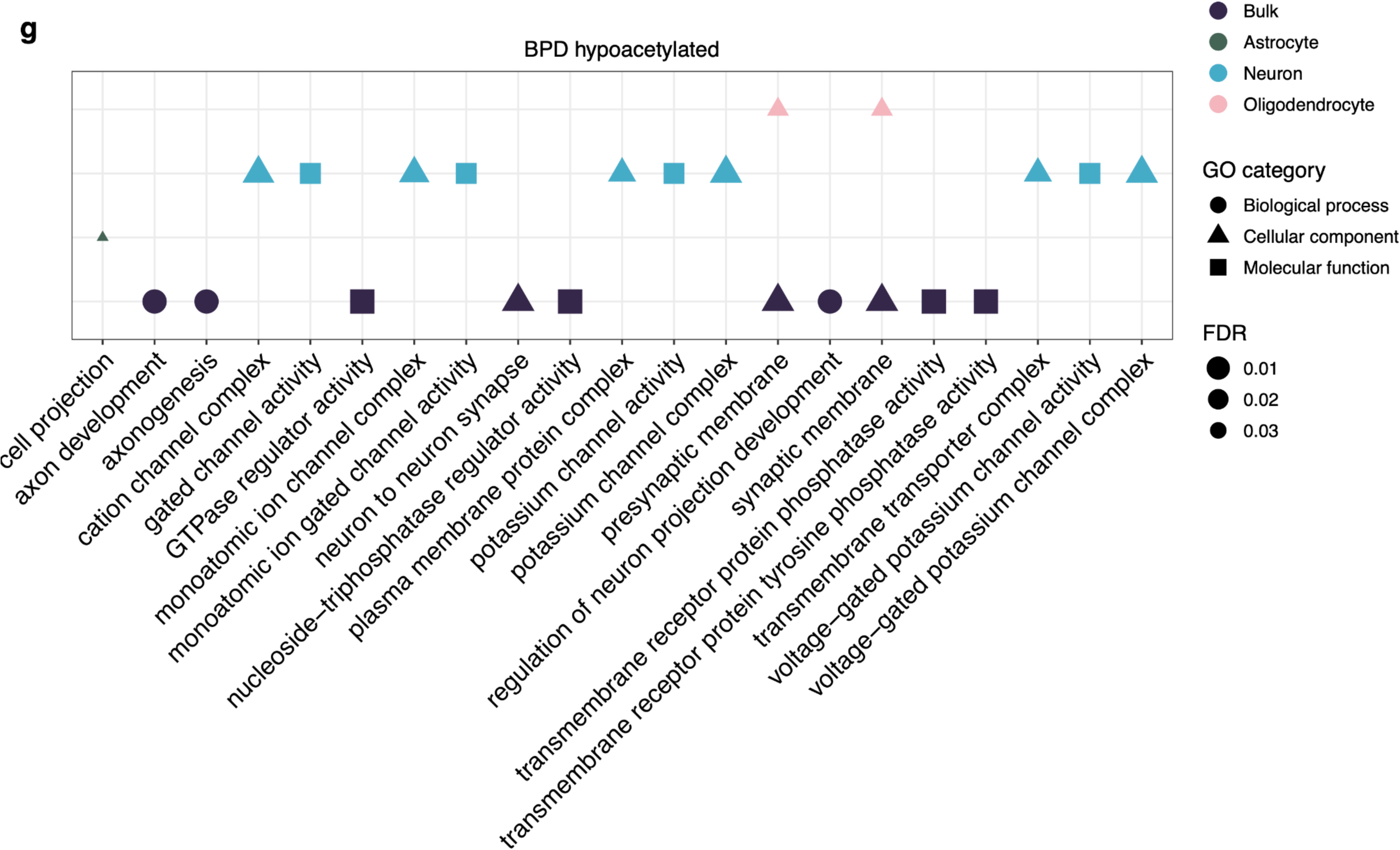
Deconvolution of bulk PFC H3K27ac in schizophrenia and bipolar disorder highlights neuron- and oligodendrocyte-specific epigenetic dysregulation. **a** Stacked barplot of proportion of bulk PFC H3K27ac peaks (schizophrenia cases, bipolar cases and controls) annotated to astrocytes, microglia, neurons, oligodendrocytes, multiple (at least two cell types), or other (none of the reference cell type peaks) using CHAS. **b** Boxplots comparing the CHAS cell type score means between schizophrenia cases and controls, and bipolar cases and controls. In both schizophrenia and bipolar cases, neurons had higher proportions and oligodendrocytes had lower proportions. Microglia had lower proportions in schizophrenia cases. P values were calculated using Welch’s two-sample t-test. **c** Stacked barplot of average cell type proportions across all bulk PFC H3K27ac profiles from schizophrenia cases, bipolar cases and controls, estimated using CHAS-MF. **d** Boxplots comparing the MF cell type proportions between schizophrenia cases and controls, and bipolar cases and controls. Changes in MF proportions when comparing cases versus controls were fairly consistent with changes in CHAS scores. P values were calculated using Welch’s two-sample t-test. **e** Heatmaps of cell type enrichments amongst schizophrenia- and bipolar-associated DARs identified when controlling for CHAS scores and CHAS-MF proportions, calculated using a. Hypergeometric p values were corrected using FDR. **f** Results from s-LDSC using summary statistics for bipolar disorder and schizophrenia GWAS and CHAS-annotated cell type-specific H3K27ac peaks from the brains of individuals with bipolar disorder and controls. In the top panel, the y-axis is the enrichment value output by LDSC. In the bottom panel, the y-axis represents the coefficient p value transformed from the coefficient z-score output by LDSC. The grey dashed line at -log10(P) = 2.4 is the cutoff for Bonferroni significance. Bipolar disorder SNP heritability was found to be exclusively enriched in neuron-specific H3K27ac peaks. **g** Functional enrichment analysis using bulk brain and cell type-specific schizophrenia-associated hypoacetylated regions. Shown are the top 10 enriched pathways for each cell type. P values were corrected using FDR.

Changes in scores and proportions when comparing cases versus controls were fairly consistent (**Fig 6c,d**). In the schizophrenia brain, we found a higher neuronal score (*p* = 0.001) and proportion (*p* = 5.5 x 10-5) when compared to the controls, and a lower microglial score (*p* = 1.6 x 10^-5^) and proportion (*p* = 1.4 x 10^-6^). In the bipolar disorder brain, we found a higher neuronal score (*p* = 0.004) and proportion (*p* = 0.0064), and a lower oligodendrocyte score (*p* = 0.019) and proportion (*p* = 0.005). Using matched samples, we compared oligodendrocyte proportion estimates derived using an independent deconvolution method in the original study, with the CHAS-derived oligodendrocyte scores, and found a significant correlation (Spearman’s rank correlation coefficient, r = 0.95, *p* < 2.2^-16^; **Supplementary Fig. 1f**).

Differential H3K27ac analysis controlling for age at death, sex, and CHAS scores revealed 235 hyperacetylated and 290 hypoacetylated peaks when comparing schizophrenia cases with controls (FDR < 0.05, **Supplementary Table 4**). When the scores were swapped for MF proportions as the covariates, this dropped down to 118 hyperacetylated and 102 hypoacetylated peaks (FDR < 0.05). Applying the same workflows to the bipolar disorder H3K27ac data, we identified 1,568 hyperacetylated and 1,298 hypoacetylated peaks when controlling for age at death, sex, and CHAS scores (FDR < 0.05, **Supplementary Table 4**). In contrast, only 159 peaks were differentially acetylated when controlling for MF-derived proportions (FDR < 0.05). Given the smaller set of disease-associated peaks identified when controlling for MF proportions, we performed cell type- and pathway enrichment analyses using the results from the differential H3K27ac analysis when controlling for CHAS scores. For both schizophrenia and bipolar disorder, DARs were most strongly enriched for oligodendrocytes (**Fig 6e**). The top ranking neuron-specific schizophrenia-associated hypoacetylated peak was located ∼1.4 kb downstream of *VGF*, which has been reported to have differential expression in schizophrenia patients^51,52^. In bipolar disorder, we identified common oligodendrocyte-specific differentially acetylated peaks in the vicinity of *AP1S2*, *SIL1*, and *EDAR*, while controlling for CHAS scores and MF proportions respectively (**Supplementary Table 4**). Overall, the differential acetylation analysis whilst controlling for cell type scores and that controlling for proportions were strongly concordant for both H3K2ac datasets (**Supplementary Fig. 2d,e**). Previous gene regulatory studies of psychiatric disorders have largely been neurocentric, whether performed using bulk tissue or neuronal populations. Our results emphasise the significance of studying cell type-specific epigenetic effects, and warrant further investigation of oligodendrocytes in bipolar disorder and schizophrenia. This is particularly relevant in schizophrenia, where transcriptomic and methylomic analyses have highlighted disease-associated changes in this cell type^21^.

Using GWAS for schizophrenia^53^ and bipolar disorder^54^, we quantified the enrichment of risk SNPs within cell type-specific peaks, and found that the heritability of both psychiatric disorders was exclusively enriched within neuronal-specific H3K27ac peaks (**Fig 6f**). This is consistent with the original study, in which the authors additionally investigated NeuN^+^ H3K27ac profiles and found that these were most strongly enriched for schizophrenia heritability^10^. Functional enrichment analysis showed that neuron-specific peaks that were hypoacetylated in bipolar disorder were associated with channel activity and complexes (**Fig. 6g; Supplementary Table 4**). While hypoacetylated peaks specific to oligodendrocytes were associated with synaptic membrane pathways (**Fig. 6g; Supplementary Table 4)**. Additional bulk and cell type-specific functional enrichments can be seen in **Supplementary Fig. 6**.

## Discussion

Given that histone acetylation is highly cell type-specific, inferring and deconvolving cell type-specific signatures in bulk tissue H3K27ac profiles is critical to our interpretation of these profiles in brain disorders. As a result, we developed CHAS, a novel tool that implements two independent algorithms for cell type deconvolution of bulk brain H3K27ac profiles: CHAS generates cell type-specific scores based on average histone acetylation levels for astrocytes, microglia, neurons, and oligodendrocytes, whereas CHAS-MF uses matrix factorisation to estimate the proportions for the same cell types. Encouragingly, when applied to bulk H3K27ac profiles from diverse brain disorders, CHAS scores and CHAS-MF proportions were highly correlated, with parallels in downstream analyses results. To the best of our knowledge, CHAS is the only publicly available tool for deconvolution of histone acetylation.

CHAS showed highly convincing performance on pseudobulk H3K27ac profiles of known cell type composition yielding near perfect correlations with the true underlying cellular proportions. These correlations remain stable at decreased sample sizes and at lower read depths, highlighting the robustness of our approach. Additionally, we were encouraged by the significant correlation between CHAS derived cell type scores and cell type proportions estimated using independent methods in the AD, bipolar disorder, and schizophrenia studies analysed here.

To illustrate the utility of CHAS for interpretation of bulk tissue H3K27ac profiles, we applied it to four epigenome-wide association studies of brain disorders^6,7,9^. Deconvolution of H3K27ac profiles from the AD brain using both CHAS algorithms highlighted that differential acetylation in late stage AD is enriched for oligodendrocyte specific H3K27ac. An independent study also found the largest H3K27ac changes in oligodendrocytes in the hippocampus and dorsolateral PFC of individuals with AD^25^. Taken together, these data suggest that these oligodendrocyte-specific H3K27ac changes are not limited to a single brain region and warrant further investigation of the role of oligodendrocyte H3K27ac dysregulation in AD. AD-associated hypoacetylation was also observed in microglia, which could reflect an increase in the activity of histone deacetylases (HDACs). In line with this, a recent study found that genetic ablation of microglial HDAC1 and HDAC2 in an AD mouse model reduced amyloid plaque burden and rescued memory deficits^55^. Thus suggesting modulation of HDAC activity in microglia as a potential therapeutic intervention for AD. Whereas epigenetic variation in late-stage AD predominantly points to oligodendrocytes, genetic risk points to microglia, suggesting independent biological mechanisms. In addition, this corroborates the finding that genetic risk for AD is enriched at microglia-specific regulatory elements^5,25,56^ and genes^36^. SImilarly, for the ASD, bipolar disorder, and schizophrenia datasets, genetic risk was enriched for neuronal-specific H3K27ac, whereas epigenetic dysregulation was most strongly associated with microglia in ASD, and oligodendrocytes in both bipolar disorder and schizophrenia. Again, the genetic risk enrichments corroborate what has previously been reported in the literature^21,42,43,57^. In the context of epigenetics in ASD, hyperacetylated promoters were linked to upregulated microglial genes^58^, and differentially methylated regions were shown to be enriched within microglial open chromatin regions^59^. In addition to highlighting which cell types are epigenetically dysregulated across diverse brain disorders, our analyses guide prioritisation of gene and pathway targets in these cell types. For example, the top AD DARs were oligodendrocyte-specific and annotated to genes involved in an oligodendrocyte gene network associated with AD. Taken together, our findings highlight the potential of CHAS in revealing biological insights and aiding prioritisation of relevant cell types and pathways for genetic and epigenetic studies of brain disorders.

There are important caveats to consider in relation to CHAS. The power of CHAS to detect cell type-specific peaks from bulk histone acetylation profiles is inherently linked to the proportion of those cells in the bulk tissue sample. This may in turn affect the robustness and power of the downstream generation of the cell type specific score and proportions, as well as the ability to detect differential acetylation in low-frequency cell types. The inclusion of additional covariates via the cell type-specific scores or proportions can contribute to a decrease in power to detect disease-associated differences in smaller sized studies. Reassuringly, controlling for cell type proportion in moderate and large sized studies, as in the case of AD and ASD, appears to improve power to detect acetylation differences, presumably via removing noise from variation caused by cellular heterogeneity. The cell sorted data on which CHAS is based has three limitations: 1) It is only available for four major brain cell types of the cortex, excluding rarer cell types such as pericytes or endothelial cells. 2) It is limited with regard to cell subtype diversity and thus unable to distinguish between functionally and regionally distinct types of neurons. 3) It is limited with regard to different cell states. For instance, multiple microglial phenotypes have been transcriptionally and functionally characterised^60–6263^. Our current reference is based on a neuropathology-free, pediatric dataset in which such states – if present – are aggregated into one category. Importantly, the performance of CHAS in bulk cerebellum samples also highlights the importance of using brain-region specific reference datasets, to account for regional differences in cell types and states. At present, CHAS is limited to bulk brain studies of H3K27ac because of reference atlas availability. However, CHAS can easily be extended to other brain cell types, regions or completely distinct tissues, as well as different histone modifications, with appropriate reference datasets. Most promisingly, we hope that future availability of single cell H3K27ac profiles across brain regions will enable us to adapt CHAS to deconvolute more refined cell subtypes and states. Finally, concerns have been raised regarding the extent to which given cell deconvolution methods can appropriately account for cellular heterogeneity, particularly when a specific trait is linked to a cellular change such as neuronal loss in AD ^16^. In this regard, it is reassuring that we observe enrichment of AD-associated epigenetic dysregulation in oligodendrocytes, but not neurons, which is in line with results reported by independent epigenomic and transcriptomic studies in human and mouse ^1464^.

Going forward epigenetic studies at single cell resolution promise to create a more comprehensive picture of the epigenetic landscape associated with neurological and psychiatric disorders. However, as single-cell H3K27ac profiling is still in its early stages of application, generates sparse data, and has not yet been achieved in the human brain, CHAS provides a unique opportunity to infer cell type-specific signatures in bulk brain histone acetylation profiles. Importantly, this can yield novel insights into epigenetic changes contributing to brain disorder risk and progression that are associated with specific cell types. Finally, CHAS is freely and publicly available at https://github.com/Marzi-lab/CHAS, adding to the existing repertoire of methods for cell type deconvolution.

## Methods

### CHAS cell type scores

CHAS is split into two independent algorithms. In the first, cell type-specific H3K27ac annotation is based on H3K27ac profiles from purified populations of astrocytes, microglia, neurons, and oligodendrocytes. These profiles were generated using H3K27ac chromatin immunoprecipitation sequencing (ChIP-seq) on 500,000 nuclei from each cell type-specific population^5^. For a given human bulk brain H3K27ac dataset, CHAS overlaps peaks detected in bulk with each cell type peak set to identify cell type-specific peaks within the bulk H3K27ac profiles. Overlaps are identified using the GenomicRanges package in R^65^. Of note, no specific threshold of overlap is required for this first cell type annotation step: even if a bulk peak overlaps a purified cell type reference peak by only one base pair, it is annotated to the cell type in this first stage. To derive the cell type-specific scores, we wanted to ensure that only high-confidence and highly cell type-specific peaks are included and therefore the following two criteria must be met: (i) the bulk peak is annotated only to a single cell type; (ii) the bulk peak overlaps a predefined percentage of the given cell type peak. This predefined percentage can be specified in the *CelltypeSpecificPeaks()* function and is set to 50% by default and for all analyses presented in this manuscript.

For each sample, CHAS generates a cell type-specific score by averaging the normalised signal intensity across all peaks specific to a given cell type. First, read counts across peaks are converted to CPM to account for variation in library size. To further normalise the signal intensity at a given peak, for each peak p, the counts are divided by the highest observed read count for that peak, thereby placing the peak-normalised counts on a scale between 0 and 1. For a sample y and cell type x, the peak-normalised counts per million s_p,y_ are summed up across all peaks p that are specific to cell type x, divided by the total number of cell type specific peaks for cell type x, P_x_. In addition, each score is also normalised by the number of cell type-specific peaks. This normalisation ensures that cell types with a greater number of specific peaks are assigned a higher CHAS score, but do not enable cross-cell type comparisons. As a constraint from peak-normalisation, the maximum signal intensity for any given peak and sample is 1 and the resulting CHAS will lie between 0 and 1 for a given sample and cell type.

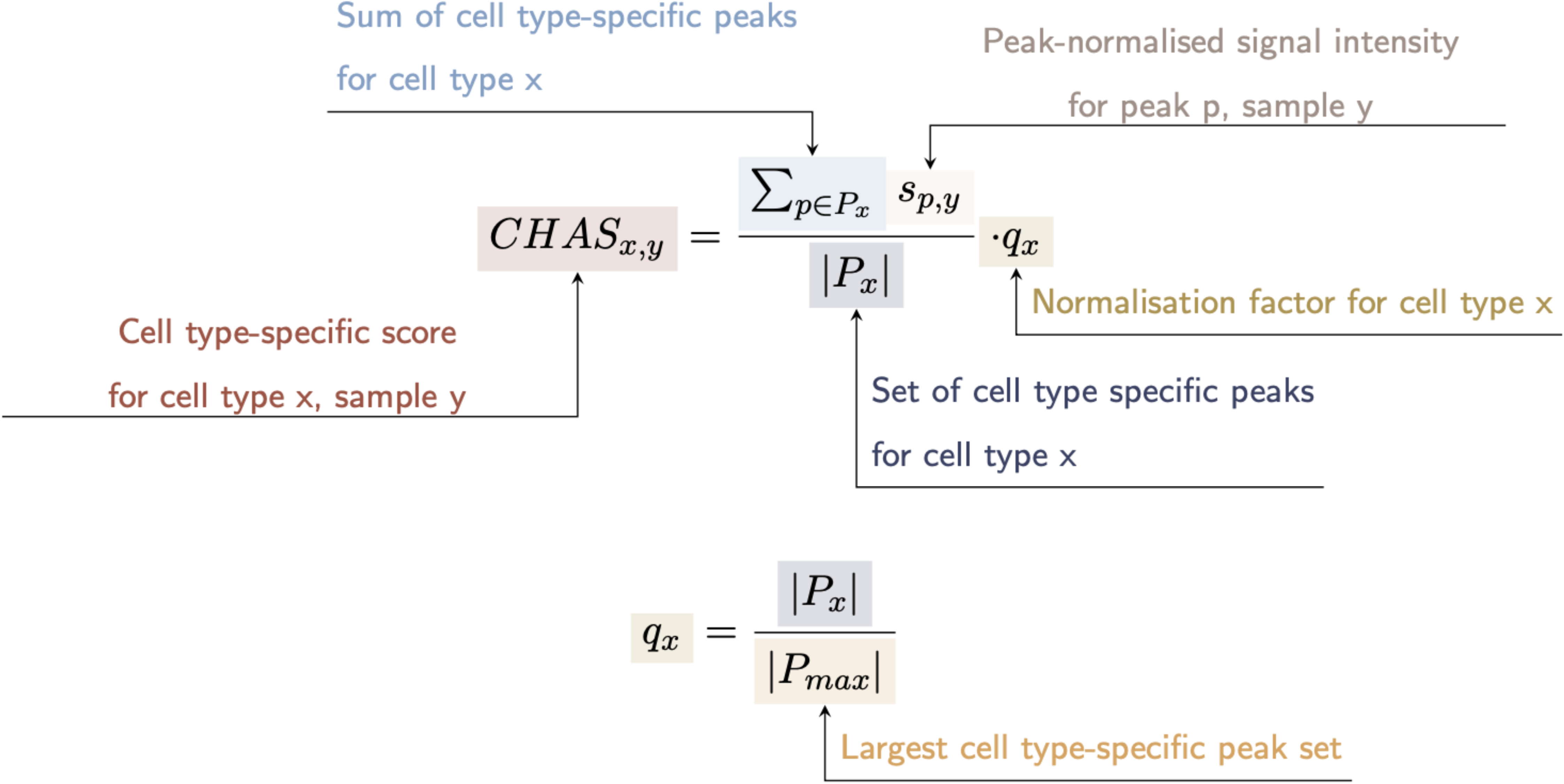

### CHAS matrix factorization

The second algorithm implemented within CHAS employs a non-negative matrix factorization approach, which is based on the R package EPIC^17^.This approach uses matrix factorisation to model bulk counts as the sum of cell type-specific counts multiplied by the corresponding cell type proportion. The bulk count matrix is modelled as the product of the reference count matrix and the cell proportion estimation matrix. EPIC solves this equation and estimates the cell proportions using constrained least square optimization^17^

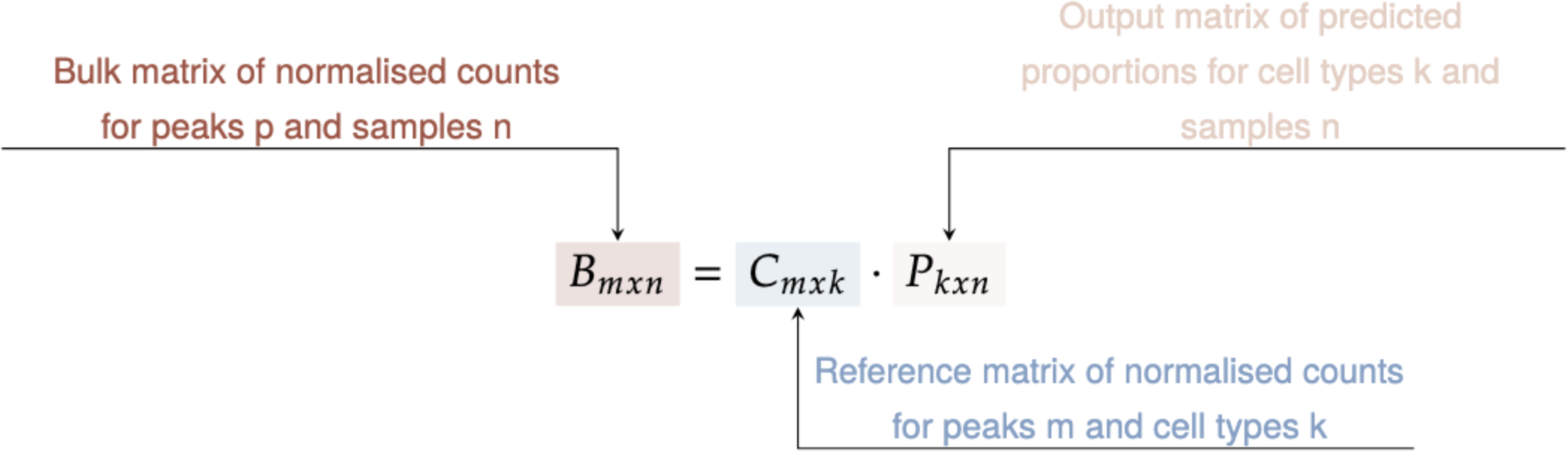

As EPIC was originally designed for transcriptomic data, we added the following steps to improve its accuracy for deconvolving bulk H3K27ac profiles. These steps are undertaken prior to running EPIC deconvolution. First, the bulk and reference H3K27ac counts are normalised based on peak length and library size to match the implicitly normalised RNA-seq counts. This is done by first dividing the raw count by the base pair length of its corresponding peak, then converting it to a CPM matrix in EdgeR. This is in direct analogy to the transcripts-per-million (TPM) normalization implemented in EPIC, modelling length and counts of H3K27ac peaks instead of transcripts (formula 2, EPIC manuscript^22^).

Second, if there are multiple reference samples for a given cell type, the median normalised CPM is used in the deconvolution. To account for signal variability within each cell type, the peaks are assigned weights for matrix factorisation. This prioritises peaks with low count variability and is derived as in the original EPIC manuscript (formula 7), using half of the range of CPM across samples as the variability metric.

Third, instead of running matrix factorisation on all consensus peaks, CHAS-MF selects a set of signature peaks with cell-type-specific signals. These are peaks that have high acetylation signals in one cell type and low signals in all the other cell types. As a threshold to qualify as signature peak, the normalised CPM for the cell type with the highest signal must be at least five times the signal from any other cell type.

The first step in CHAS-MF is merging bulk peaks and reference peaks to create consensus peaks - which are defined as the union of bulk and reference peaks. Each consensus peak is then annotated with corresponding cell types, depending on whether the consensus peak overlaps cell type-specific peaks. Read counts are generated for the consensus peaks, either from bam files or existing counts for the bulk and reference samples. Using this count data, CHAS-MF predicts the proportion of each cell type in bulk samples by applying the EPIC R package, using our histone acetylation optimized adaptations. The calculation is performed on normalised read counts for signature peaks, controlling for library size and peak lengths. If multiple reference samples are used for one cell type, the median normalised CPM will be used as the main input for matrix factorisation, and the signal variability for each peak is taken into account as a weight term in the matrix factorisation.

The cell type annotation, generation of cell type specific histone acetylation scores, and generation of MF-derived cell type proportion estimates are implemented and automated in our R package CHAS (https://github.com/Marzi-lab/CHAS).

### Validation of CHAS

We validated CHAS using three independent approaches. First, by simulating pseudobulk H3K27ac profiles of known cell type composition based on the raw sequencing data from astrocytes, microglia, neurons and oligodendrocytes by Nott and colleagues^5^. The cell type composition of each pseudobulk sample was based on proportions of astrocytes, microglia, neurons, oligodendrocytes and endothelial cells in brain tissue from older individuals, which had been quantified in an independent study using immunohistochemistry^23^. Based on the reported cell type proportions of 49 post-mortem brain samples we generated 49 pseudobulk samples, pooling a total of 30 million randomly sampled reads per sample from the raw H3K27ac data of the four cell types. As our reference did not include H3K27ac profiles for endothelial cells, we excluded the proportion of this cell type and instead used the relative proportions of the four other cell types. We ran CHAS and CHAS-MF to generate cell type-specific scores and cell type proportions, respectively, for each pseudobulk sample and compared these to the true cell type proportions using Pearson correlation coefficients. To additionally evaluate the robustness of CHAS with respect to sample size and sequencing depth, this process was repeated across the 49 samples with 20 million and 10 million randomly sub-sampled reads, as well as using 30 million reads in random subsets of 25 and 10 samples. Second, we used CHAS with an existing dataset of NeuN^+^ and NeuN^-^ H3K27ac peaks from the anterior cingulate cortex (n=15) and dorsolateral PFC (n=14). The *CelltypeSpecificPeaks()* function was used to annotate each peak set, and then using the peaks that were annotatable to at least one cell type, we calculated the CHAS scores for each cell type. We then used a paired t-test to compare the average neuronal proportion across the NeuN^+^ peaks with that of the NeuN^-^ peaks. As the CHAS-MF workflow requires H3K27ac read counts, the associated NeuN^+^ and NeuN^-^ bam files were downloaded and pre-processed in the same way as was done for the AD H3K27ac dataset (see Methods: H3K27ac in entorhinal cortex from AD cases and controls (Marzi et al. 2018)). Consensus peaks were identified using the *ConsensusPeaks()* function on the processed peaks and counts data. The cell typr proportion was estimated using the *CelltypeProportion()* function. We repeated the t-test comparing the average MF-derived neuronal proportion of NeuN^+^ peaks with that of the NeuN^-^ peaks. Finally, for the AD^7^ and schizophrenia and bipolar disorder^10^ datasets, we were able to correlate CHAS-derived neuronal and oligodendrocyte scores, respectively, with cellular proportions quantified independently in the original studies. In the AD H3K27ac study, neuronal proportions were quantified based on methylation profiles using CETS^11^, and in the schizophrenia and bipolar disorder H3K27ac study, oligodendrocyte proportions were quantified based on acetylation profiles using dtangle^66^.

## Application of CHAS to bulk brain H3K27ac datasets

### H3K27ac in entorhinal cortex from AD cases and controls (Marzi et al. 2018)

In order to demonstrate reproducibility and undertake preprocessing using updated versions of software and the most recent reference genome, raw ChIP-seq data from our previous study was downloaded from sequence read archive (SRA) under accession number PRJNCA297982^7^. We performed basic quality control using fastQC^67^. Using bowtie2^50^ the fastq files were aligned to the most recent human reference genome (GRCh38)^68^. The resulting SAM files were converted to binary (BAM) format using SAMtools^69^. Duplicates, unmapped reads, and reads with a sequence quality score q < 30 were removed from all BAM files and the filtered BAM files were subsequently merged into one grouped file. Next, using MACS2^39^ we performed peak calling on the merged file of all samples. The following peak sets were subsequently filtered out: 1) peaks which overlapped the ENCODE blacklist peaks (https://github.com/Boyle-Lab/Blacklist), 2) peaks which were located in unmapped contigs or mitochondrial DNA, and 3) peaks which did not meet a significance threshold of *P* < 10^-7^ for peak calling. Read count generation for each sample was performed using featureCounts70 and read counts were converted to and stored in a CPM matrix, keeping peaks with a minimum of three samples showing ≥ 1 read per million. This resulted in a total of 183,353 peaks to be used in downstream analyses. This optimal peak set and CPM matrix were used as input to run CHAS to identify cell type-specific peaks in the bulk H3K27ac profiles and to generate cell type-specific H3K27ac scores as a proxy for the proportion of each cell type in the bulk peak set. Of note, to annotate cell types to each bulk peak and to calculate the cell type proportions across the bulk peaks, we only required an overlap of at least one base pair between the bulk peak and the cell type peak. However, for peaks included in the cell type specific histone acetylation score we required a more stringent overlap of at least 50% of the cell type peak interval. We also ran CHAS-MF on the processed peaks and counts along with the BAM files to estimate the proportions of each cell type in the bulk samples. The CHAS-generated cell type-specific scores and CHAS-MF-derived proportions were used to detect shifts in cellular composition between AD cases and controls, by comparing the means using a Welch’s t-test. Differences in histone acetylation between AD cases and controls were analysed as previously described^7^, but including the CHAS derived cell type scores or MF proportions, instead of the neuronal proportion estimator based on CETS^11^. Briefly, the quasi-likelihood F test in the Bioconductor package edgeR^24^ was used to test for differences in histone acetylation between AD cases and controls, while controlling for: (i) age at death and cell type-specific scores for the four brain cell types; (ii) age at death and cell type proportions. All covariates were treated as continuous numeric variables. Peaks were considered differentially acetylated at FDR<0.05. To additionally confirm that we had adequately controlled for false-positive associations, we permuted the AD case and control labels 100 times and repeated the differential histone acetylation analysis as described above.

### H3K27ac in PFC from PD cases and controls: Toker et al. 2021

Peak lists and read count tables for the Park West (PW) cohort were downloaded from https://github.com/ltoker/ChIPseqPD. Peak lists were in narrowPeak format and were filtered to include peaks mapping to canonical chromosomes, and to exclude peaks which overlapped those in blacklisted regions https://github.com/Boyle-Lab/Blacklist, as well as those not meeting a significance threshold of P < 10-7 for peak calling. For the PW cohort, a total of 171,285 peaks were used for downstream analyses. From the counts tables we excluded the sample outliers identified in Toker et al (2021) and performed final filtering, keeping peaks with a minimum of three samples showing ≥ 1 read per million for the differential histone acetylation analysis. This left us with 152,823 peaks in the PW cohort. The counts table along with the filtered peak set were used as input to CHAS and CHAS-MF, as previously described for the AD dataset. As there were no BAM files available for the PD study, we performed CHAS-MF using the original counts for the bulk and reference peaks as a proxy for the read counts for consensus peaks. The CHAS-generated cell type-specific scores and CHAS-MF-derived proportions were used to detect shifts in cellular composition between PD cases and controls, by comparing the means using a Welch’s t-test. Differences in histone acetylation between PD cases and controls were analysed as described above, controlling for age at death, sex, and cell type proportions using the: (i) CHASscores, or (ii) CHAS-MF proportions.

### H3K27ac in PFC and cerebellum from ASD cases and controls: Sun et al. 2016

ChIP-seq reads mapped to the human reference genome (hg19) using BWA^71^ by Sun and colleagues (2016) were downloaded from Synapse under accession number syn4587616. We downloaded 80 libraries from the PFC and 62 libraries from the cerebellum. These were the same libraries that were used in the original study for peak calling^6^, with exception of one PFC sample which was not available on Synapse.

Downloaded files were in BAM format and all pre-processing steps were performed as described previously (see H3K27ac in entorhinal cortex from AD cases and controls: Marzi et al. 2018 under Methods). The optimal peak sets for downstream analyses totaled 250,614 peaks for PFC, and 241,759 peaks for cerebellum. These, alongside the counts matrices, were used as input to CHAS. We also ran CHAS-MF on the same peaks and counts data, along with the BAM files, to estimate the proportion of each cell type in the bulk samples. Differences in histone acetylation between ASD cases and controls for each brain region were analysed as described above, controlling for age at death, sex, and: (i)CHAS-derived cell type scores, (ii) CHAS-MF proportions. Disease-associated differences in cell type proportions were quantified using a Welch’s t-test on the CHAS-derived cell type-specific scores and the CHAS-MF derived cell type proportions.

### H3K27ac in PFC from schizophrenia and bipolar disorder cases and controls: Girdhar et al. 2022

Raw sequencing data (FASTQ files) for 249 samples were downloaded from Synapse under accession number syn25705564. Of these libraries, 68 were from schizophrenia cases, 48 were from bipolar disorder cases, and 133 were from controls. All pre-processing, mapping, and peak calling steps were performed as described previously (see H3K27ac in entorhinal cortex from AD cases and controls: Marzi et al. 2018, under Methods), with peaks being called on a merged file containing all schizophrenia, bipolar disorder, and control samples. 396,065 peaks were used as input to CHAS and CHAS-MF, as well as for differential H3K27ac analysis controlling for sex, age at death, and: (i) cell type-specific histone acetylation scores, (ii) cell type proportions. Disease-associated differences in cell type proportions were quantified using a Welch’s t-test on the CHAS-derived cell type-specific scores and the CHAS-MF derived cell type proportions.

### Quantifying enrichments of cell type-specific H3K27ac peaks within disease-associated differentially acetylated regions

To test whether the proportion of cell type-specific peaks in the disease-associated DARs differed significantly from the background set of cell type-specific non-DARs, we used the hypergeometric test. P values were corrected using the FDR. This was calculated for the AD, ASD, bipolar disorder, and schizophrenia H3K27ac peaks.

### Genomic annotation and enrichment analysis

Gene annotation and gene ontology analyses were performed using clusterProfiler^28^. Gene annotation was performed using the *annotatePeak()* function using the default parameters and additionally specifying *annoDb="org.Hs.eg.db"* to retrieve gene symbols. Gene ontology analysis was performed using the *enrichGO()* function, for the ontology categories biological process, molecular function, and cellular component. Enrichment P values were corrected for multiple testing using an FDR cutoff of 0.05.

### Partitioned heritability analysis

To estimate the proportion of disease SNP-heritability attributable to cell type-specific H3K27ac peaks identified in bulk brain data, we performed partitioned heritability analysis as implemented in LDSC^34^. For each cell type-specific peak set, annotation files were generated and used to compute LD scores. Publicly available GWAS summary statistics for an AD GWAS^35^, ASD GWAS^72^, PD GWAS^38^, schizophrenia GWAS^53^, and bipolar disorder GWAS^54^, were downloaded and converted to the required format for LDSC. Steps for the analysis were followed as instructed here https://github.com/bulik/ldsc/wiki. For each annotation, LDSC was run using the full baseline model^34^, thereby computing the proportion of SNP-heritability associated with the annotation of interest, while taking into account all the annotations in the baseline model.

### Data and code availability

CHAS is a free and publicly available R package: https://github.com/Marzi-Lab/CHAS. All the data and code required to reproduce the figures in this manuscript are available in our GitHub repository: https://github.com/Marzi-lab/CHAS_manuscript. All supplementary tables are available at: 10.5281/zenodo.12784761.

## Funding

SJM and AN are supported by the Edmond and Lily Safra Early Career Fellowship Program and the UK Dementia Research Institute [award number UKDRI-6009 and UKDRI-5016] through UK DRI Ltd, principally funded by the Medical Research Council. SJM received funding from the Alzheimer’s Association (grant number ADSF-21-829660-C) and the MRC (grant number MR/W004984/1). KBM was funded by the UK Medical Research Council Doctoral Training Partnership (https://mrc.ukri.org).

## Author contributions

Conceptualization: SJM

Methodology: SJM

Software: KBM, YY

Formal Analysis: KBM, YY

Validation: KBM, YY

Investigation: AN, SJM

Data Curation: KBM, SJM, AN, MT

Writing – original draft: KBM, SJM

Writing – review & editing: KBM, YY, SJM, AN, MT

Visualization: KBM, YY

Supervision: SJM

Project administration: SJM

## Acknowledgments

We thank Alan E Murphy and Brian M Schilder at the UK Dementia Research Institute, Imperial College London for feedback and helpful discussions on the CHAS R package.

## Supplementary information

**Supplementary Figure 1:**
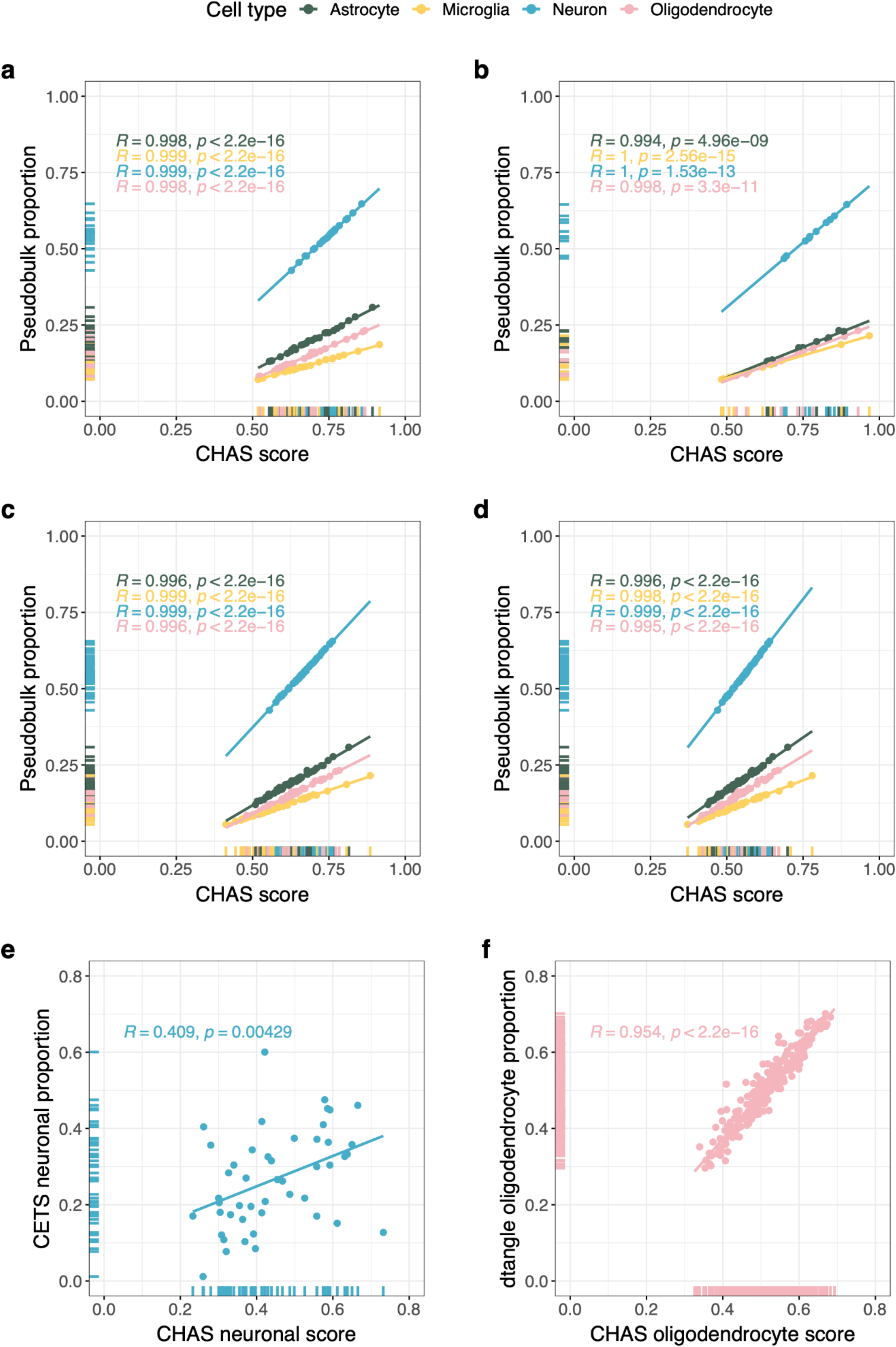
CHAS robustly estimates cell type proportions in datasets with low coverage and small sample size, and correlates with known cell type proportion. **a-d** Validation of CHAS using pseudobulk samples made up of randomly sampled reads from astrocytes, microglia, neurons, and oligodendrocytes. Shown are scatterplots of the CHAS-derived histone acetylation score (x-axis) vs. the true proportion of the cell type within the pseudobulk sample (y-axis) for **(a)** 25 samples with 30 million reads each, **(b)** 10 samples with 30 million reads each, **(c)** 49 samples with 20 million reads each, and **(d)** 49 samples with 10 million reads each. Pearson correlation coefficient R and p value are shown for each individual cell type. CHAS score had near perfect correlation with the actual cell proportion in all scenarios. e-f Validation of CHAS using neuronal proportion (NeuN⁺ fraction) estimates derived using CETS^11^ in the AD study^7^ and oligodendrocyte proportion estimates derived using dtangle^66^ in the schizophrenia and bipolar study^10^. **(e)** CHAS-derived neuronal scores and CETS-derived neuronal proportion estimates had a significant correlation (R = 0.41, p = 0.004) across the 47 samples from AD patients and controls. CETS neuronal proportion estimates in this sample were derived based on DNA methylation profiles from the same entorhinal cortex tissue block. **(f)** CHAS-derived oligodendrocyte scores and dtangle-derived oligodendrocyte proportion estimates had a significant correlation (R = 0.95, p < 2.2 x 10^16^) across the 249 samples from the schizophrenia and bipolar patients and controls.

**Supplementary Figure 2:**
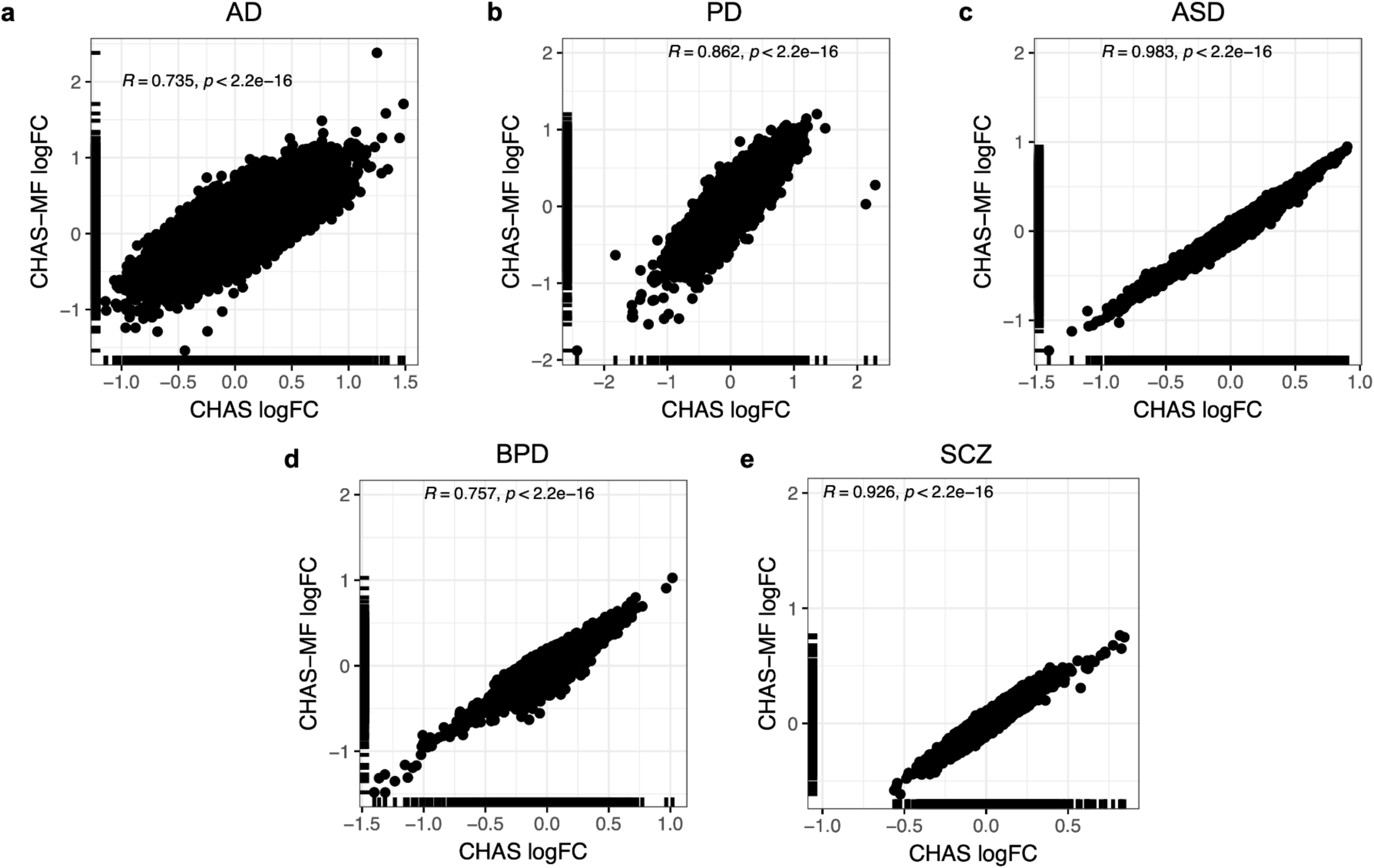
Acetylation changes quantified whilst controlling for cell type scores and CHAS-MF derived cell type proportions correlate strongly. Scatterplots of logFC when performing differential acetylation analysis whilst controlling for CHAS-derived scores vs CHAS-MF-derived cell type proportions in the a AD H3K27ac dataset^7^, b PD H3K27ac dataset^9^, c ASD H3K27ac dataset^6^, d,e BPD and SCZ H3K27ac dataset^10^.

**Supplementary Figure 3.**
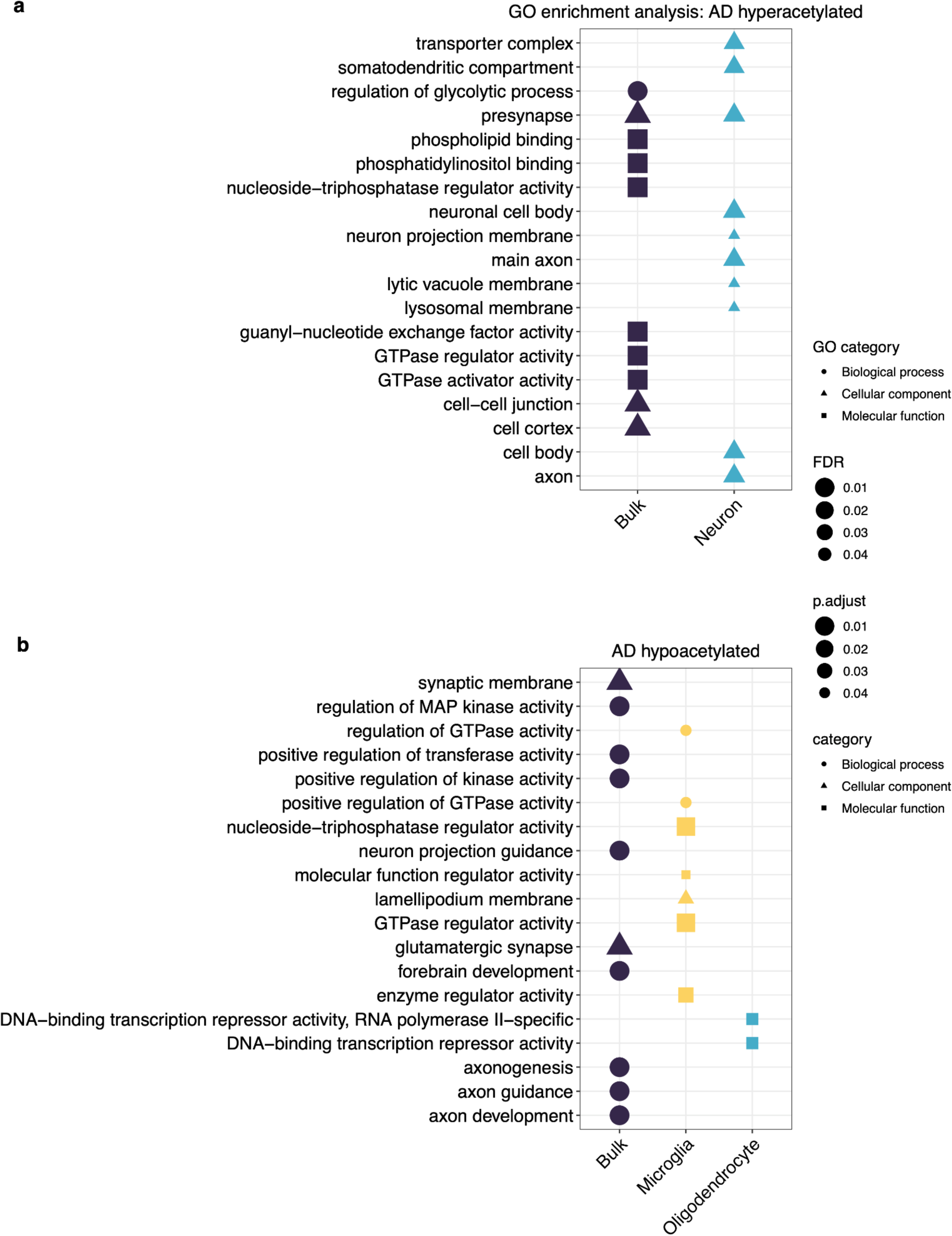
Functional enrichment analysis using AD-associated H3K27ac regions. Pathway enrichment analysis using AD-associated bulk and cell type-specific a hyperacetylated peaks controlling for CHAS scores, (**b**) hyperacetylated peaks controlling for MF-derived proportions, and (**c**) hypoacetylated peaks controlling for MF-derived proportions. Shown are the top 10 enriched pathways for each cell type. P values were corrected using FDR.

**Supplementary Figure 4.**
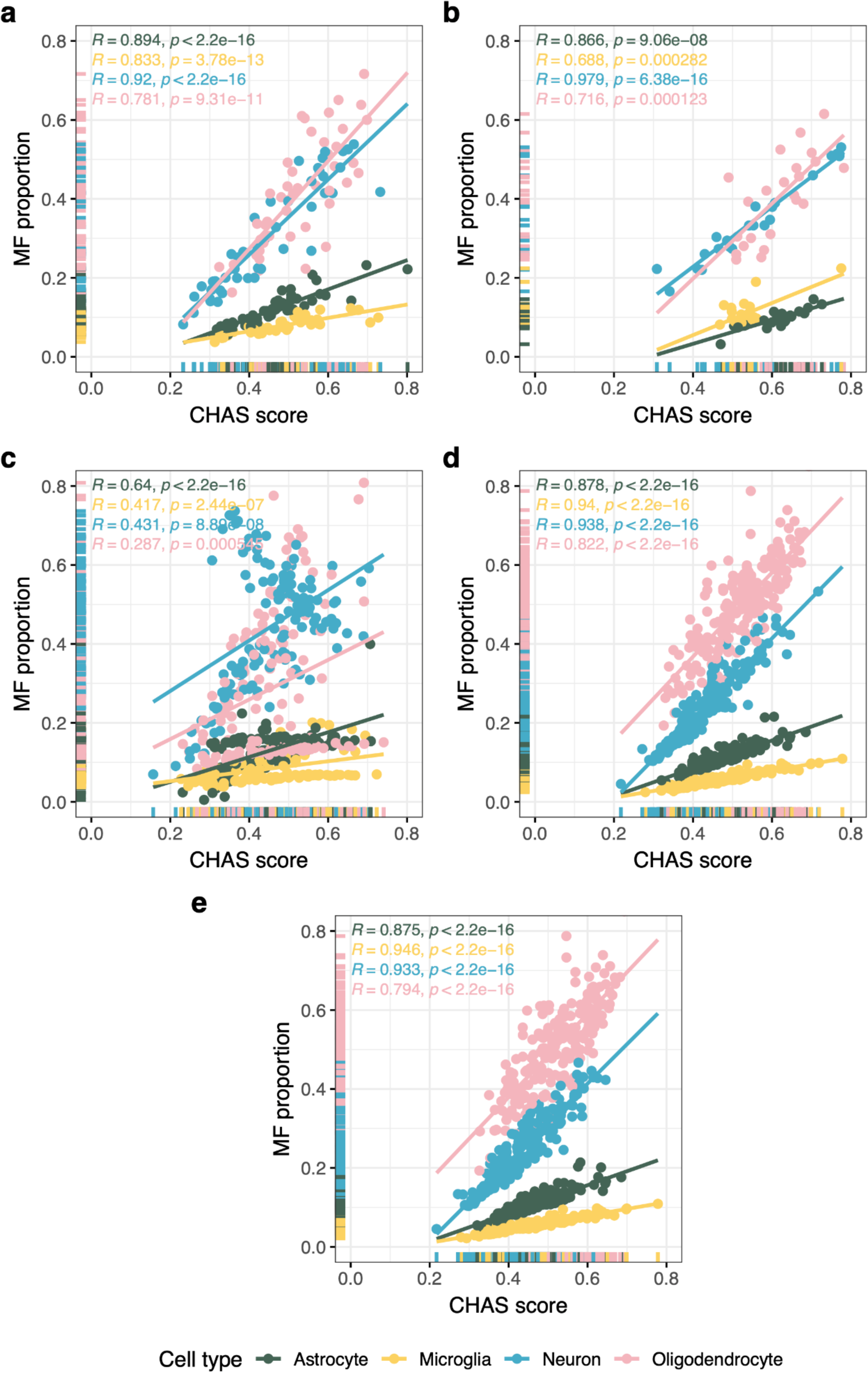
Correlations of CHAS scores and CHAS-MF proportions for each brain disorder H3K27ac dataset. **a** Scatterplot of CHAS-derived scores vs. CHAS-MF-derived proportions for the AD H3K27ac dataset. **b** Scatterplot of CHAS-derived scores vs. CHAS-MF-derived proportions for the PD H3K27ac dataset. **c** Scatterplot of CHAS-derived scores vs. CHAS-MF-derived proportions for the ASD H3K27ac dataset. **d** Scatterplot of CHAS-derived scores vs. CHAS-MF-derived proportions for the BPD H3K27ac dataset. **e** Scatterplot of CHAS-derived scores vs. CHAS-MF-derived proportions for the SCZ H3K27ac dataset. Spearman’s rank correlation coefficient *r* and *p* values are shown for each cell type.

**Supplementary Figure 5.**
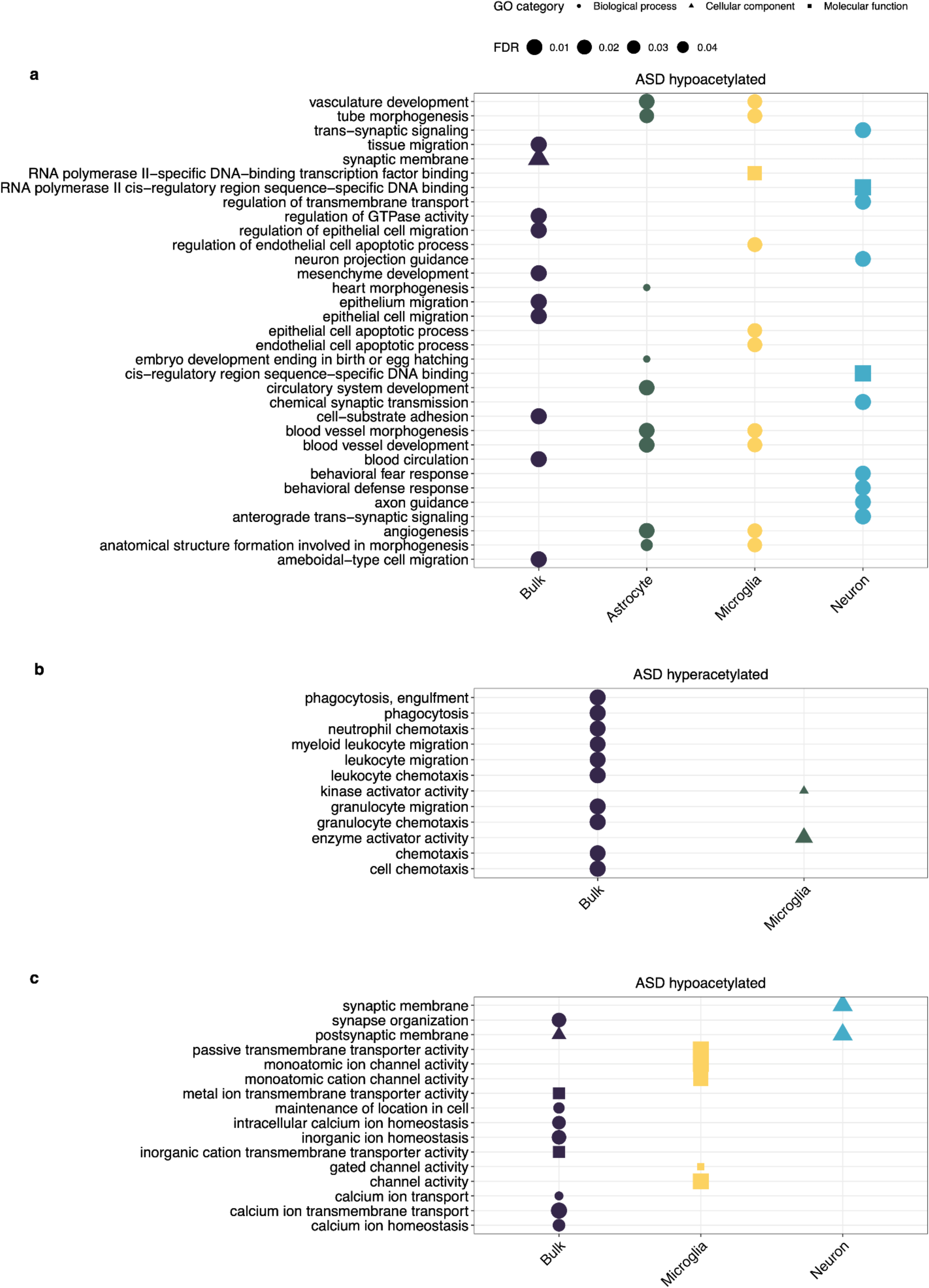
Functional enrichment analysis using ASD-associated H3K27ac regions. GO enrichment analysis using ASD-associated bulk and cell type-specific **a** hypoacetylated peaks controlling for CHAS scores, **b** hyperacetylated peaks controlling for MF-derived proportions, and **c** hypoacetylated peaks controlling for MF-derived proportions. Shown are the top 10 enriched pathways for each cell type. P values were corrected using FDR.

**Supplementary Figure 6.**
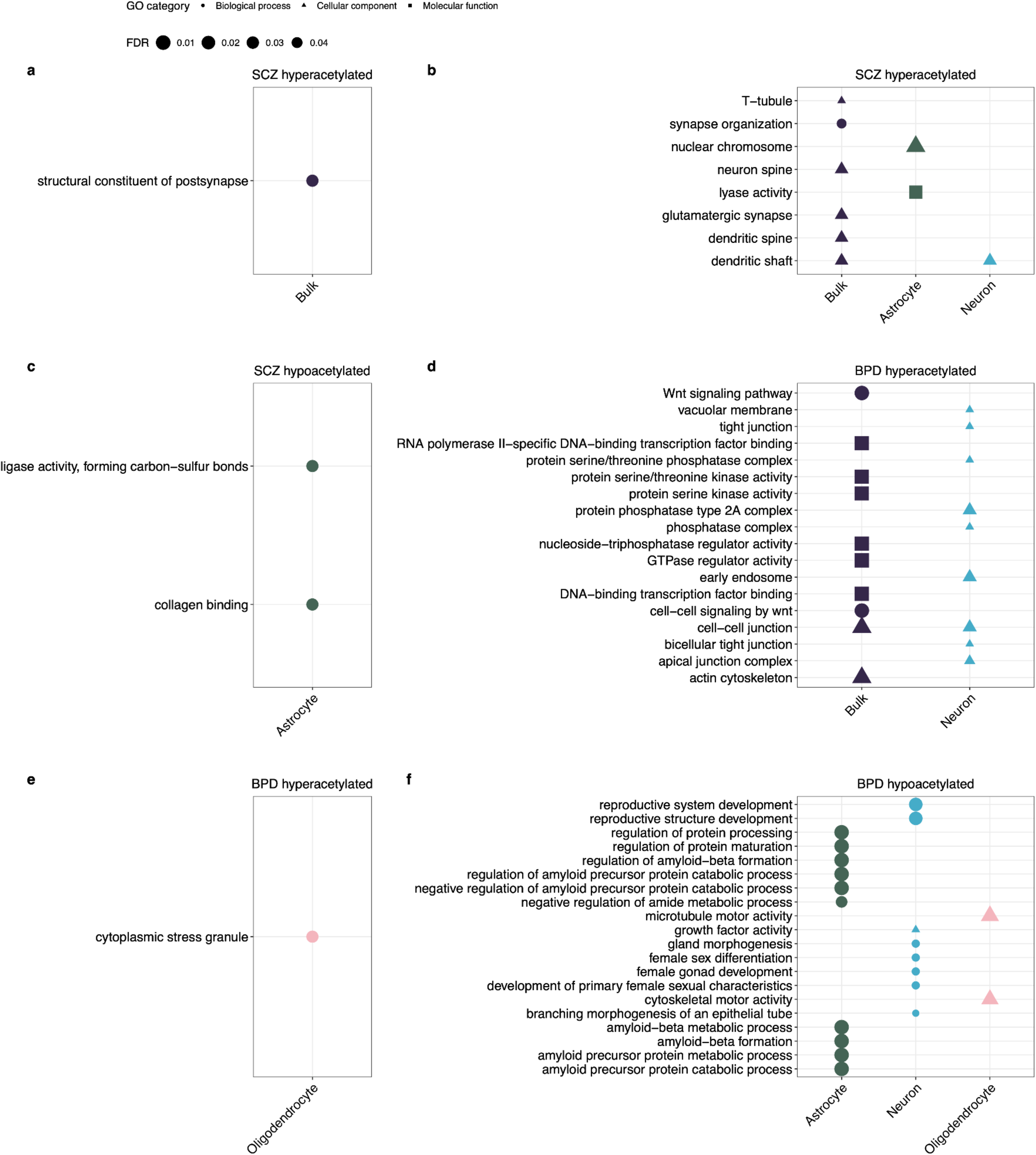
Functional enrichment analysis using schizophrenia-associated and bipolar disorder-associated H3K27ac regions. **a-c** Pathway enrichment analysis using schizophrenia-associated bulk and cell type-specific a hyperacetylated peaks controlling for CHAS scores, (b) hyperacetylated peaks controlling for MF-derived proportions, and (c) hypoacetylated peaks controlling for MF-derived proportions. d-f Pathway enrichment analysis using bipolar disorder-associated bulk and cell type-specific (d) hyperacetylated peaks controlling for CHAS scores, (e) hyperacetylated peaks controlling for MF-derived proportions, and (f) hypoacetylated peaks controlling for MF-derived proportions. Each plot shows the top 10 enriched pathways for each cell type. P values were corrected using FDR.

## Notes

https://github.com/Marzi-lab/CHAS

https://github.com/Marzi-lab/CHAS_manuscript

https://doi.org/10.5281/zenodo.12784761

### Competing Interest Statement

The authors have declared no competing interest.

### Summary of Updates

Cell type identity is a major driver of epigenetic variation, making the biological interpretation of bulk tissue epigenomes challenging. Here, we present CHAS (Cell Type-Specific Histone Acetylation Score), an R package for the cell deconvolution of bulk brain H3K27ac profiles, which allows us to control for cellular heterogeneity and infer cell type-specific epigenetic patterns associated with neurological and psychiatric disorders. This manuscript includes the following major updates 1) We implemented a second, independent algorithm for cell type proportion estimation based on non-negative matrix factorisation 2) We added additional validation, showing that CHAS can accurately distinguish neuronal and non-neuronal cell types and correlates with other cell type proportion metrics. 3) We additionally applied CHAS to a large post-mortem brain study of schizophrenia and bipolar disorder.

